# Efficient in vivo neuronal genome editing in the mouse brain using nanocapsules containing CRISPR-Cas9 ribonucleoproteins

**DOI:** 10.1101/2022.07.24.501299

**Authors:** Jeanette M. Metzger, Yuyuan Wang, Samuel S. Neuman, Kathy J. Snow, Stephen A. Murray, Cathleen M. Lutz, Viktoriya Bondarenko, Jesi Felton, Kirstan Gimse, Ruosen Xie, Yi Zhao, Matthew T. Flowers, Heather A. Simmons, Subhojit Roy, Krishanu Saha, Jon Levine, Marina E. Emborg, Shaoqin Gong

**Affiliations:** Wisconsin National Primate Research Center, University of Wisconsin-Madison, Madison, WI, 53715; Department of Ophthalmology and Visual Sciences, University of Wisconsin–Madison, Madison, WI, 53715; Department of Biomedical Engineering, University of Wisconsin–Madison, Madison, WI, 53715; Wisconsin Institute for Discovery, University of Wisconsin–Madison, Madison, WI, 53715; The Jackson Laboratory, Bar Harbor, ME, 04609; Departments of Pathology and Neuroscience, University of California-San Diego, San Diego, CA, 92093; Department of Neuroscience, University of Wisconsin–Madison, Madison, WI, 53715; Department of Medical Physics, University of Wisconsin-Madison, Madison, WI, 53715

**Author notes:** Corresponding authors;, Dr. Shaoqin Gong: Wisconsin Institute For Discovery, 330 North Orchard Street Room 4166, Madison WI 53715, 1-608-316-4311, Dr. Marina Emborg: Wisconsin National Primate Research Center 1220 Capitol Court, Madison WI 53715, 1-608-262-9714. These two authors contributed equally and are listed alphabetically (Jeanette M. Metzger, Yuyuan Wang).

## Abstract

Genome editing of somatic cells via clustered regularly interspaced short palindromic repeats (CRISPR) offers promise for new therapeutics to treat a variety of genetic disorders, including neurological diseases. However, the dense and complex parenchyma of the brain and the post-mitotic state of neurons make efficient genome editing challenging. *In vivo* delivery systems for CRISPR-Cas proteins and single guide RNA (sgRNA) include both viral vectors and non-viral strategies, each presenting different advantages and disadvantages for clinical application. We developed non-viral and biodegradable PEGylated nanocapsules (NCs) that deliver preassembled Cas9-sgRNA ribonucleoproteins (RNPs). Here, we show that the RNP NCs led to robust genome editing in neurons following intracerebral injection into the mouse striatum. Genome editing was predominantly observed in medium spiny neurons (>80%), with occasional editing in cholinergic, calretinin, and parvalbumin interneurons. Glial activation was minimal and was localized along the needle tract. Our results demonstrate that the RNP NCs are capable of safe and efficient neuronal genome editing *in vivo*.

**SIGNIFICANCE STATEMENT:** Modifying the DNA of cells in the brain could present opportunities for new treatments of neurological diseases. In this report, we describe a nanocapsule system designed to deliver the elements needed to modify the DNA of brain cells, also known as genome editing. These nanocapsules are created by chemically encapsulating the genome editing components, such that the nanocapsules are stable when prepared and biodegradable to release their payload upon entering cells. When injected into the mouse brain, our research shows that the nanocapsules lead to safe and efficient editing of DNA in neurons.

## INTRODUCTION

CRISPR-Cas9 *in vivo* editing of somatic cells holds significant promise for treating rare and common diseases ^1,2^. RNA-guided Cas9 systems can quickly and efficiently cleave target DNA in coding or non-coding areas of the genome with low off-target effects. CRISPR-Cas9 genome editing has been tested in multiple animal species as a method of generating disease models and as a potential therapy. The technology has recently moved into clinical trials to treat several pathologies including cancer ^3^, hereditary transthyretin amyloidosis ^4^, and an inherited cause of childhood blindness ^5^. Building on these advances, newer technologies capable of safely delivering CRISPR-Cas9 genome editors to the brain and inducing robust neuron editing could revolutionize the treatment of neurological disorders ^6,7^.

Efficient genome editing of the neurons in the central nervous system (CNS) presents notable challenges. The vasculature of the CNS forms the blood brain barrier (BBB), which controls brain homeostasis by tightly regulating the movement of ions, molecules, and cells between the bloodstream and the brain. For systemic administration, compounds targeting the CNS need to have specific properties to cross the BBB^8^. High dosages of these compounds may be required that could produce unwanted side effects ^9^. Temporary disruption of the BBB is proposed as an alternative approach for BBB penetration, but this approach carries unique risks associated with negating the protection of the BBB ^9,10^. Intracerebral injection, albeit invasive, allows for bypassing the BBB by direct delivery into the brain parenchyma. Regardless of the route or delivery method, once inside the brain, the delivered substance must then traverse a dense and complex neuropil to access neurons. Particles with smaller sizes and neutral charges are advantageous for brain editing, as they can diffuse over longer distances in this unique extracellular matrix ^11-13^.

Viral or plasmid vectors have been successfully applied for *in vivo* delivery of Cas9 and single guide RNA (sgRNA) to the brain, most commonly through the use of adeno-associated virus (AAV) vectors ^14-18^. To deliver genome editing components, AAVs depend on the host transcriptional and translational machinery of the cell to generate genome editing ribonucleoproteins (RNPs). Further, their genetic DNA payload is limited by the packaging ability of AAV (∼5kb) ^19^. These vectors are typically considered to present relatively low risk for integration into the genome; yet, recent work has demonstrated that AAV vectors carrying CRISPR components frequently integrate at the site of double strand breaks ^20,21^. The resultant prolonged expression of the Cas9 nuclease may increase the risk for eventual off-target effects ^21-23^. Viral vector capsids and accessory proteins may trigger an immune response, impacting efficacy and biosafety *in vivo* ^24^. Antibodies against different AAV serotypes have been identified in the human population, posing a particular challenge to clinical translation ^25^. Cas9 can alternatively be introduced by nonviral delivery of mRNA, which can be impacted by low RNA stability ^23^. Moreover, lentiviral vectors to deliver Cas9 are genome integrating vectors; thus, they carry some risk of insertional mutagenesis and genotoxicity ^20^. In comparison, delivery of preassembled Cas9 protein/sgRNA RNP avoids genome integration and leads to transient Cas9 expression in the cell, thereby lowering the risks of deleterious insertional effects and off-target editing ^19,26-28^.

The first demonstration of *in vivo* brain editing using Cas9 protein/sgRNA RNP was reported in 2017 via intracerebral injection of RNPs tagged with multiple nuclear localization signals ^29^. Following this landmark study, a limited number of non-viral vectors for RNP delivery for neuronal editing have been reported, such as CRISPR-gold ^30^ and peptide nanocomplexes ^31^. We developed a novel RNP nanocapsule (NC) for RNP delivery in which monomers with different functional moieties bind to the RNP surface and form a covalently crosslinked, yet degradable, polymeric coating via *in situ* free radical polymerization ^32^. The RNP NCs achieve endosomal escape via imidazole-containing monomers and lead to release of RNP into cellular cytosol by cleavage of the glutathione-responsive cross-linker. The versatile surface chemistry of the PEGylated NCs allows for convenient conjugation of various types of targeting ligands or cell penetrating peptides (CPPs). Furthermore, in contrast to self-assembled nanoparticles, the RNP NCs have outstanding *in vivo* stability before entering the target cells, due to their covalent nature. The RNP NCs also have a much smaller size (around 35 nm) compared with other types of self-assembled nanoparticles (typically larger than 100 nm), which may facilitate their diffusion within the brain. Finally, the RNP NCs also enable a relatively high RNP loading content (∼40 wt%). The NC editing efficiency has been previously demonstrated in murine retinal pigment epithelium tissue and skeletal muscle ^32^. The aim of this study was to assess the application of these uniquely engineered RNP NCs for neuronal genome editing in the mouse brain.

## RESULTS

The RNP NCs were prepared as previously reported with minor modifications (Fig. 1a) ^32^. For intracerebral injection, we hypothesized that conjugation of a neuron-specific ligand (e.g., rabies virus glycoprotein-derived peptide, aka RVG), or a CPP (i.e., TAT peptide), can enhance the specificity for neuron-targeted delivery and/or the cellular uptake of the NCs. To test this hypothesis, acrylate-PEG-RVG and acrylate-PEG-CPP were first synthesized via a Michael addition reaction between acrylate-PEG-maleimide and thiolated RVG (or CPP) peptides. Monomers with different functional moieties (i.e., positive/negative charges, imidazole groups for endosomal escape, and acrylate-PEG with or without ligands), as well as the disulfide-containing crosslinker were coated onto the RNP surface with optimized molar ratios ^32^. After coating, free radical polymerization was initiated by the addition of ammonium persulfate and tetramethylethylenediamine. Acrylate-PEG with or without ligand was added in the last step to form PEGylated RNP NCs. The hydrodynamic diameters and zeta-potentials of NCs, as determined by dynamic light scattering (DLS), were approximately 35 nm and 2 mV, respectively, and were similar across NC formulations (i.e., NC-No Ligand, NC-RVG, and NC-CPP), as shown in Fig. 1b. NCs with different surface modifications also showed similar morphologies according to transmission electron microscope (TEM) images (Fig. 1c). The stability of NC was evaluated functionally by loading the NC with Cas9 and a sgRNA targeting green fluorescent protein (GFP) in a transgenic human embryonic kidney (HEK 293) cell line. Successful delivery of RNP by NCs results in GFP gene disruption, thus, the genome-editing efficiencies of NCs stored for different durations were evaluated by the percentage of GFP-negative cells. NC-No Ligand delivering the RNP targeting the GFP gene was prepared using sgRNAs purchased from two different companies (i.e., sgRNA 1 and sgRNA 2), and dispersed in the storage buffer. NCs were stored at different temperatures (i.e., 4°C, -20°C and -80°C), and gene editing efficiency was studied at designated timepoints. As shown in Figure 1d, NCs were stable for at least 130 days at -80°C without significant gene editing efficiency change.

**Figure 1.**
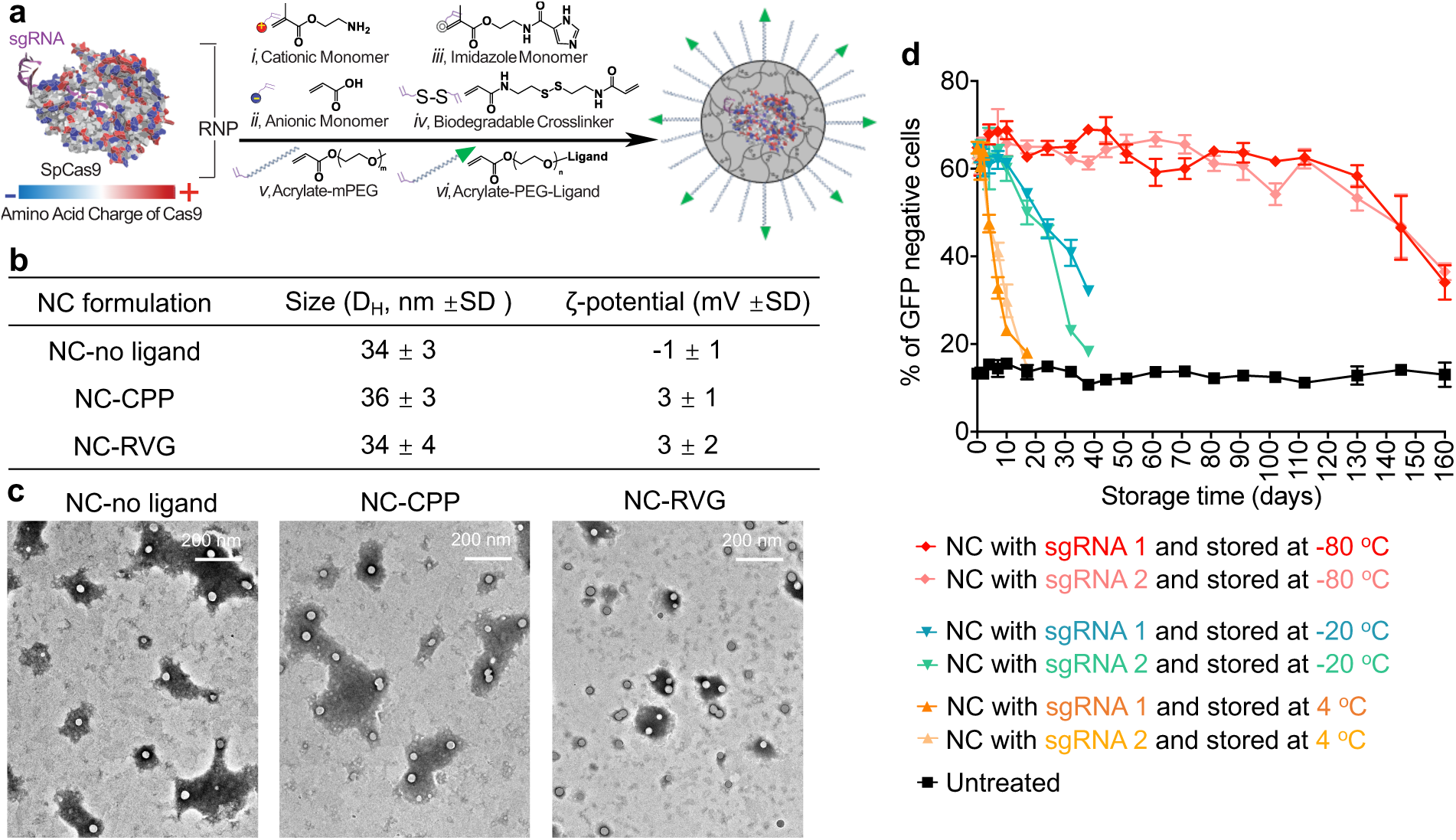
Synthesis and characterization of the RNP NCs. a, A schematic illustration for the synthesis of RNP-encapsulated NC. b, Summary of the sizes and zeta-potentials of NCs with or without ligand. c, TEM images of NC-No Ligand, NC-CPP and NC-RVG. d, RNP delivery of NC after storage at different conditions. CPP, cell penetrating peptide; GFP, green fluorescent protein; NC, nanocapsule; RNP, ribonucleoprotein; RVG, rabies virus glycoprotein; sgRNA, short guide RNA; TEM, transmission electron microscopy.

To evaluate the ability of the NCs to deliver RNP and produce *in vivo* neuronal genome editing, NCs were stereotactically injected into the striatum of Ai14 mice (Fig. 2b). The Ai14 reporter mouse harbors a LoxP-flanked stop cassette containing three SV40 polyA transcriptional terminators, which act to prevent the expression of the red fluorescent protein tdTomato. RNP-targeting of sequences within this stop cassette can lead to the expression of tdTomato when at least two SV40 polyA blocks are excised; therefore, genome editing is detectable via red fluorescence, although the fluorescent tdTomato protein underreports the total genome editing outcomes (Fig. 2a)^29,32,33^. Two weeks following stereotactic, intrastriatal NC injection, the animals were euthanized by trans-cardiac perfusion, and the brains were collected. The coronal sections across the striatum were analyzed for tdTomato positive (i.e., genome-edited) cells and co-stained for neuronal (neuronal nuclear protein, NeuN) and astroglial (glial fibrillary acidic protein, GFAP) markers (Fig. 2d-g). Regions of interest (ROIs) were defined by the extent of cells showing red florescence (Fig. 2e). These ROIs were evaluated to determine the neuronal editing efficiency and size of the genome-edited brain area (Fig. 2f,g). Based on successful development and application of these techniques in a methods development animal cohort (Supp.Fig. 1; Supp. Table 1), a larger, independent, second site (The Jackson Laboratory) study was performed to validate the results and compare the three formulations, namely, – NC-No Ligand, NC-CPP, and NC-RVG (Fig. 3; Supp. Table 1). This multi-site approach is a key feature of the National Institutes of Health (NIH) Somatic Cell Genome Editing (SCGE) consortium to ensure data reproducibility and scientific rigor.

**Figure 2.**
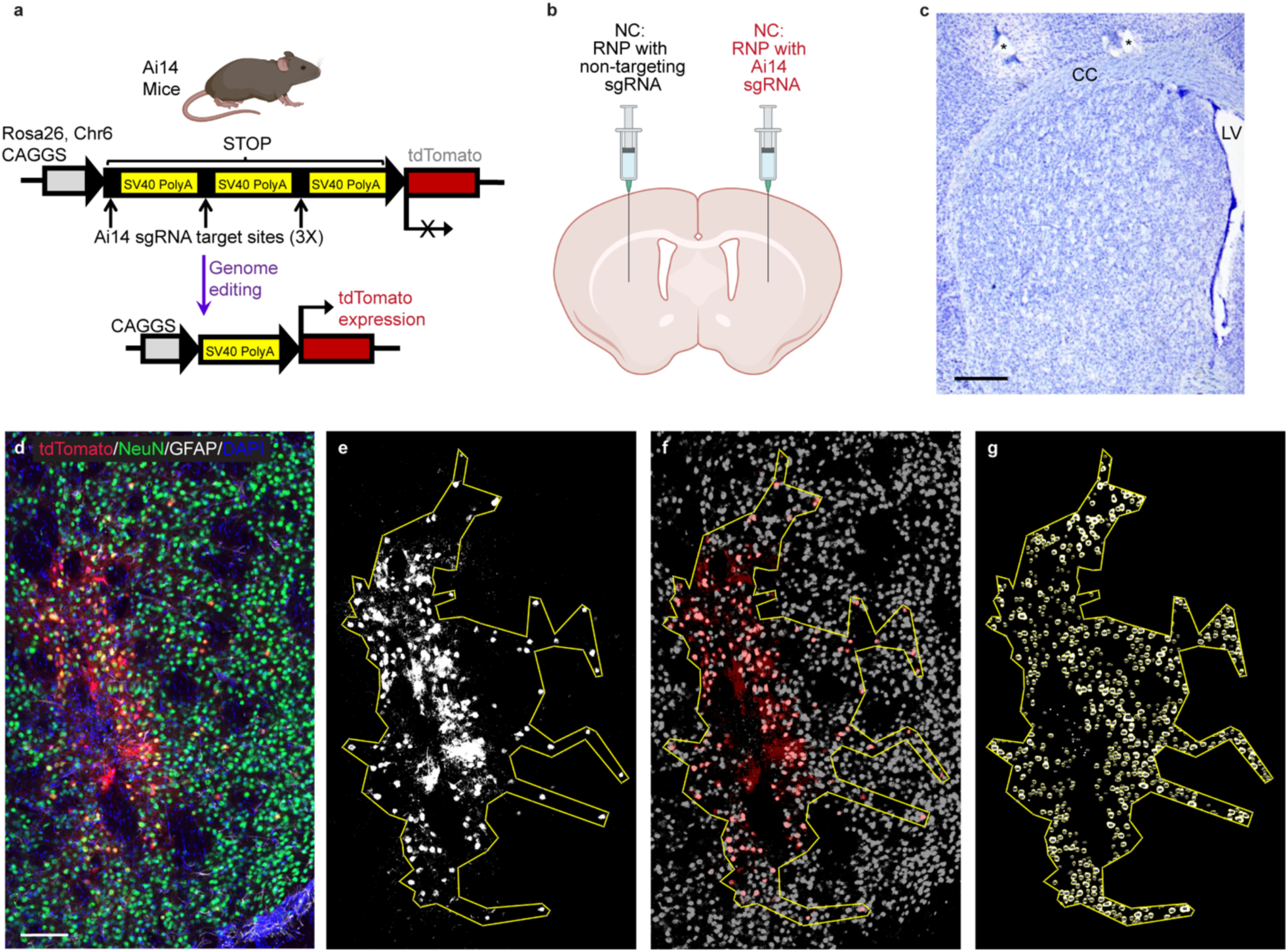
Methods used for analysis of neuronal genome editing. a, Schematic of the Ai14 mouse tdTomato locus. CRISPR/Cas9 targeting sites are present in each of the three SV40 polyA sequences in the STOP cassette. Removal of two of the SV40 polyA cassettes leads to the expression of tdTomato. b, Schematic of the intrastriatal injection of NCs. c, Nissl-stained coronal mouse section showing normal striatal anatomy. *, holes made during tissue processing to identify left hemisphere. CC, corpus callosum. LV, lateral ventricle. Scale = 500um. d – g, Steps performed for the analysis of edited area size and percent neuronal editing shown in a single representative coronal brain section (animal J4). d, Photomicrograph of genome-edited neurons in the mouse striatum showing maximum intensity projection of three focal planes covering 10 μm. Neurons (NeuN, green) that are genome-edited are tdTomato (red) expressing. GFAP (astrocytes) = white. DAPI (nuclei) = blue. Scale = 100 μm. e, Using FIJI software, a binary mask of the tdTomato channel was used to draw a region of interest (ROI) around the edited, tdTomato+ cells. f, the tdTomato binary mask (red) was then overlayed onto a NeuN signal binary mask (grey) to manually count genome-edited neurons (tdTomato+ and NeuN+). g, Using StarDist2D object identification followed by watershed segmentation of the NeuN signal, the NeuN channel was processed to allow for automated counting of the total number of neurons in the ROI. DAPI, 4′,6-diamidino-2-phenylindole; GFAP, glial fibrillary acidic protein; NeuN, neuronal nuclear protein.

**Figure 3.**
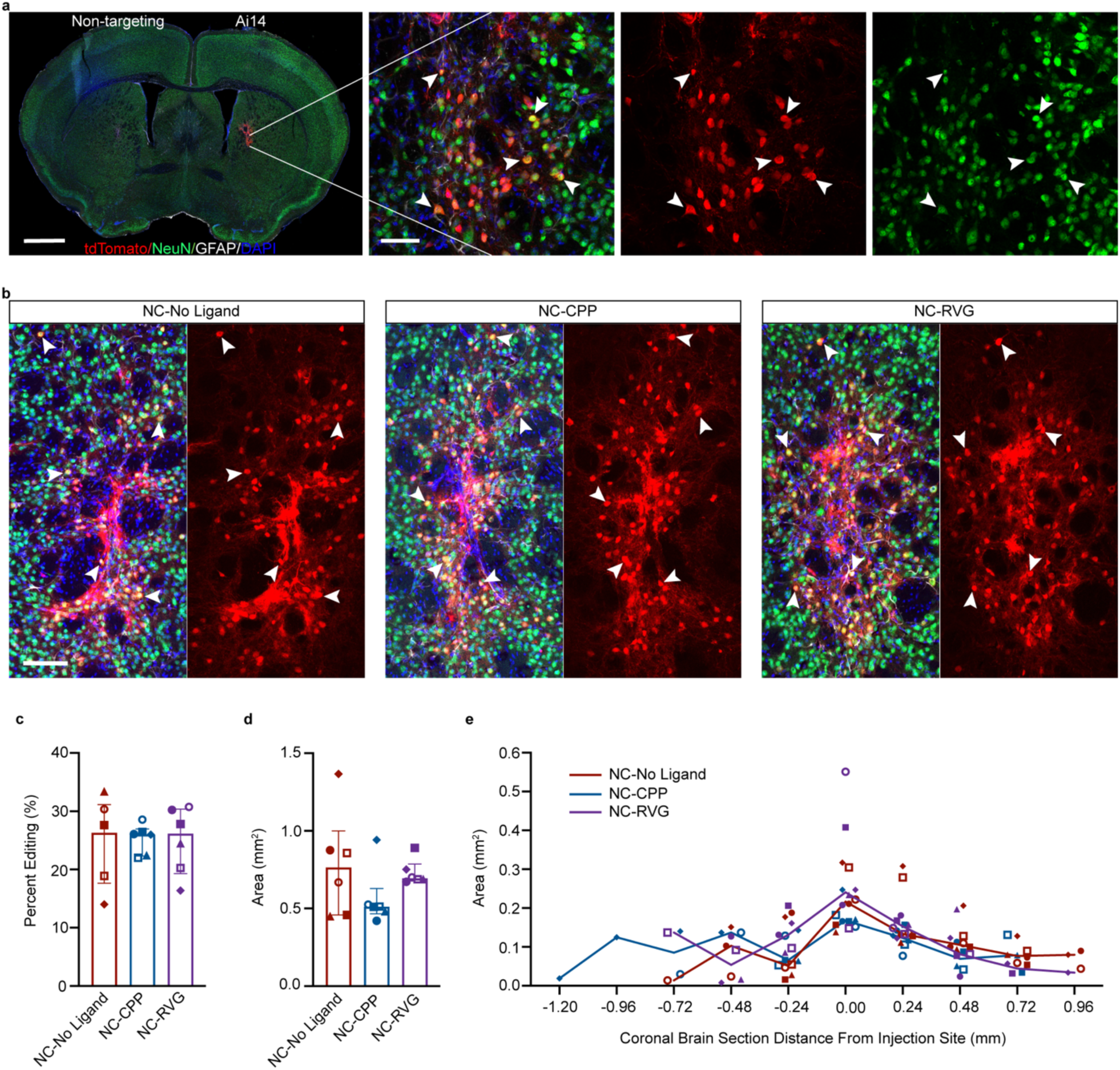
NC delivery of CRISPR RNP produces efficient *in vivo* neuronal genome editing in the striatum of Ai14 mice. a, Coronal mouse brain section (scale = 1000 μm) and high magnification image (scale = 50 μm) of neuronal genome editing following NC injection (animal J11). Co-labeling (yellow; white arrowheads) of the neuronal maker NeuN (green) and tdTomato (red) indicates genome-edited neurons. b, Genome-edited neurons (yellow; white arrowheads) in the striatum of representative animals from each of the NC treatment groups (NC-No Ligand animal J12; NC-CPP animal J18; NC-RVG animal J5; Supp. Table 1). Scale bar = 100 μm. c, Percentage of neurons in the edited area expressing tdTomato in each NC treatment group. d, Sum of edited area size (region of interest area) across all coronal slices in each NC treatment group. e, Line graph of median edited area size for each treatment group at given distances rostral and caudal to the injection site. c and d, Graphs show median and interquartile range. Differences between groups were not statistically significant. c – e, Each animal is shown with a unique color and symbol. Photomicrographs show maximum intensity projection of three focal planes covering 10 μm. Individual channels were adjusted for brightness as needed (Supp. Table 4). CPP, cell penetrating peptide; DAPI, 4′,6-diamidino-2-phenylindole; GFAP, glial fibrillary acidic protein; NC, nanocapsule; NeuN, neuronal nuclear protein; RVG, rabies virus glycoprotein.

Mice treated with NCs loaded with RNP containing the Ai14-targeting sgRNA showed successful genome editing of striatal neurons (Fig. 3a,b; Supp.Fig. 1; Supp. Table 1). At the site of injection of NCs with Ai14 sgRNA, abundant genome-edited, neuron-like cells were present, characterized by triangular cell bodies filled with intense red tdTomato fluorescence and lighter fluorescence in extensions, consistent with axons and dendrites (Fig. 3a,b). A small number of cells suggestive of astrocytes based on their polygonal shape with multiple processes also expressed tdTomato. To confirm the identity of the genome-edited cell types, triple immunofluorescence staining with antibodies against tdTomato, the astrocyte marker GFAP, and the neuron marker NeuN was performed (Fig. 3a,b). Colocalization of nearly all tdTomato signal with NeuN+ cells confirmed that the majority of the edited cells were neurons (Fig. 3).

Quantification of the neuronal editing efficiency, or the percentage of neurons (NeuN+ cells) that were genome-edited (tdTomato+ cells), indicated similar genome-editing capability across all three NC formulation groups, with 26.3% ± 9.3% in the NC-No Ligand group, 26.0% ± 3.0% in the NC-CPP group, and 26.2% ± 8.3% in the NC-RVG group (*H* = 0.035; *df* = 2; *p* = 0.990; Fig. 3c). The total genome-edited brain area, defined as the sum of the ROI areas across all analyzed coronal sections, was 0.763 mm^2^ ± 0.360 in the NC-No Ligand group, 0.512 mm^2^ ± 0.032 in the NC-CPP group, and 0.695 mm^2^ ± 0.051 in the NC-RVG (Fig. 3d). Differences between the three NC formulation groups were not statistically significant (*H* = 2.889; *df* = 2; *p* = 0.248). Rostrocaudal spread of genome-editing covered a range of approximately 1.44 mm in the NC-No Ligand and NC-CPP groups and 1.20 mm in the NC-RVG group (Fig. 3e). Comparison between cohorts demonstrated a relationship between injected volume of NCs and edited area. The edited area was significantly larger in the methods development animals receiving the 1.5 µl injections (1.474 mm^2^ ± 0.517) compared to the definitive study that were injected with 1 µl (0.680 mm^2^ ± 0.319; *U* = 11; *n*_*1*_ = 7; *n*_*2*_ = 18; *p* = 0.0008; *r* = 0.629) when comparing animals in all treatment groups of these cohorts. This effect remained statistically significant when a potential outlier (Supp. Table 1 UW3 with a total ROI size of 7.399 mm^2^) was removed from the 1.5 µl injected animal dataset (1.367 mm^2^ ± 0.510; *U* = 11; *n*_*1*_ = 6; *n*_*2*_ = 18; *p* = 0.0025; *r* = 0.585).

The majority of the edited striatal neurons were medium spiny neurons identified by co-labeling of tdTomato and dopamine- and cAMP-regulated phosphoprotein 32 kDa (DARPP32) (Fig. 4a-c, g; Supp.Fig. 2). DARPP32+ neurons accounted for approximately 80.8% ± 12.2% of all tdTomato+ neurons across all NC treatment groups, without significant differences between groups (NC-No Ligand 84.0% ± 14.4; NC-CPP 79.7% ± 17.6%; NC-RVG 79.2% ± 8.8; *H* = 1.867; *df* = 2; *p* = 0.439; Fig. 4g). Occasionally, other edited neuronal subtypes were observed, such as cholinergic (choline acetyltransferase, ChAT+), parvalbumin+, and calretinin+ neurons (Fig. 4; Supp.Figs. 2 & 3).

**Figure 4.**
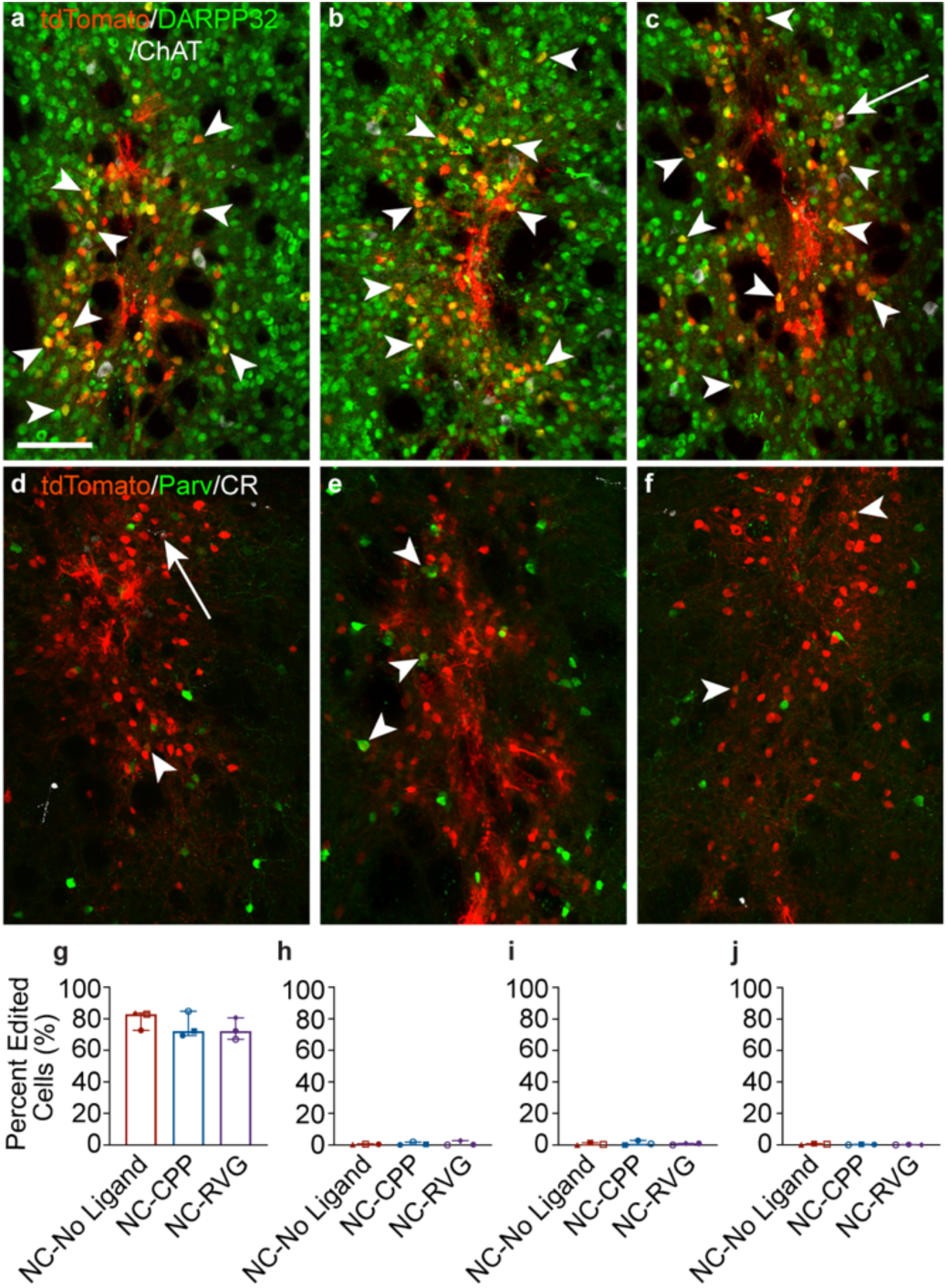
Striatal genome editing following RNP NC delivery is observed primarily in medium spiny neurons. a – c, Genome-edited neurons in the mouse striatum (tdTomato+; red) are primarily medium spiny neurons, as indicated by DARPP32 (green) co-labeling (white arrowheads). A small number choline acetyl transferase (ChAT)+ (white) neurons are genome-edited (white arrows). This pattern was similar across NC treatment groups (a, NC-No Ligand animal J9; b, NC-CPP animal J17; NC-RVG animal J5; Supp. Table 1). d – f, Genome-edited parvalbumin (Parv+; white arrowheads) and calretinin (CR+; white arrows) neurons were occasionally observed (a, NC-No Ligand animal J8; b, NC-CPP animal J13; NC-RVG animal J4; Supp. Table 1). a – f, scale = 100 μm. Photomicrographs show maximum intensity projection of three focal planes covering 10 μm. Individual channels were adjusted for brightness as needed (Supp. Table 4). Percentage of tdTomato+ neurons that co-labeled for DARPP32 (g), ChAT (h), parvalbumin (i), or calretinin (j) across treatment groups. Graphs shows median and interquartile range. Differences between groups were not statistically significant. Each animal is shown with a unique color and symbol (Supp. Table 1). CPP, cell penetrating peptide; DAPI, 4′,6-diamidino-2-phenylindole; DARPP32, dopamine- and cyclic-AMP-regulated phosphoprotein of molecular weight 32 kDa; NC, nanocapsule; RNP, ribonucleoprotein; RVG, rabies virus glycoprotein.

The host immune reaction to the RNP NC treatment was assessed in hematoxylin and eosin (HE)-labeled brain sections by a board-certified veterinary pathologist blinded to treatment groups (Fig. 5a-c). Analyses of the three treatment groups - NC-No Ligand with RNP containing Ai14 sgRNA, NC-No Ligand with RNP containing non-targeting sgRNA, and uninjected control - did not detect significant pathology. Small areas of increased cellularity were found in the cortex and striatum of hemispheres injected with both non-targeting sgRNA NCs and Ai14 sgRNA NCs in coronal tissue samples, consistent with the locations of the injections in the striatum and mechanical passage of the needle (needle tract) through the cortex.

**Figure 5.**
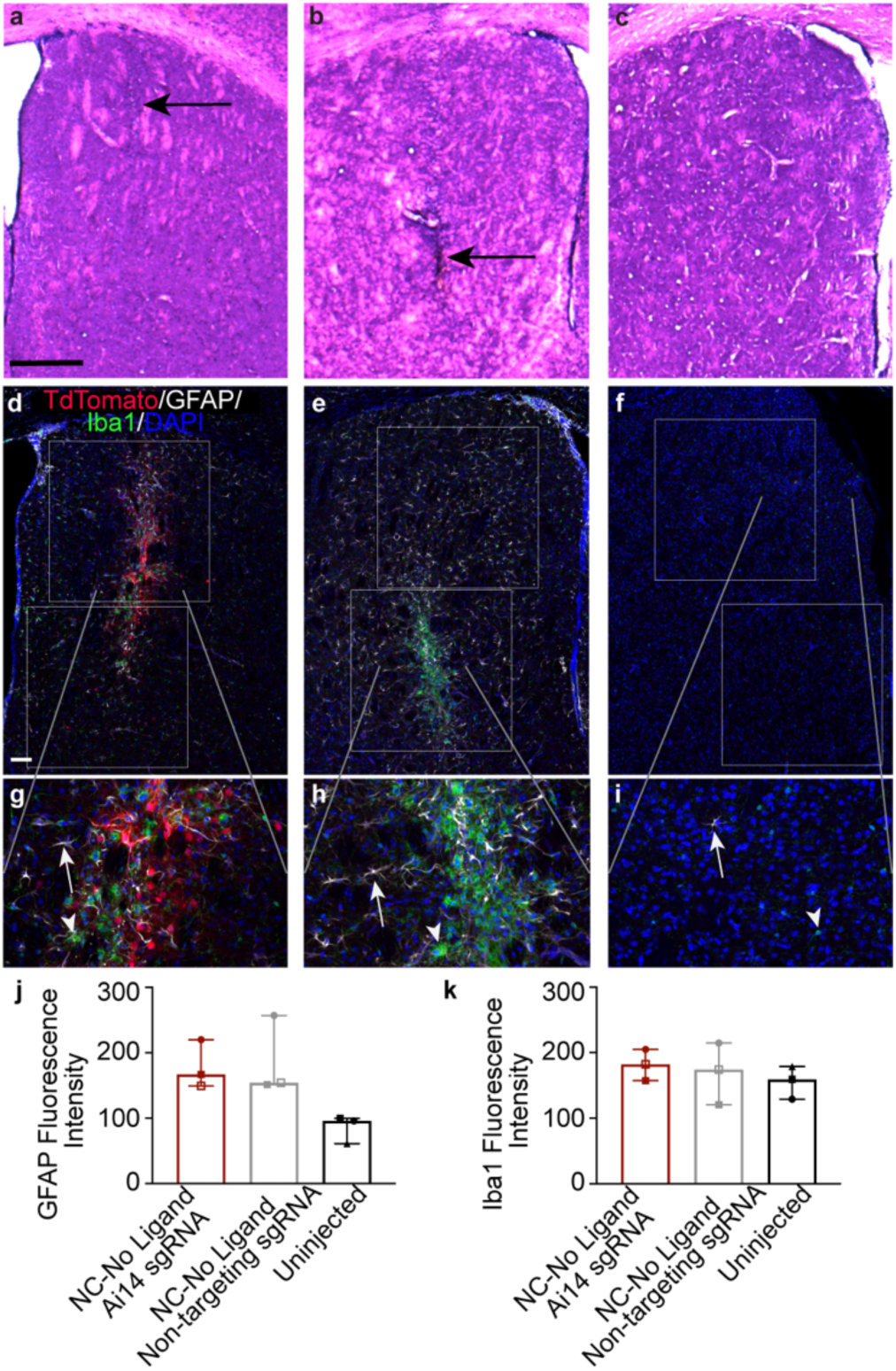
NC injection is not associated with a significant inflammatory response. a – c, Hematoxylin and eosin (HE) stained mouse brain tissue at the injection site in the striatum showing linear focal areas of increased cellularity (black arrows) in each of the treatment groups (a, NC-No Ligand RNP with non-targeting sgRNA [animal J7 left hemisphere]; b, NC-No Ligand RNP with Ai14 sgRNA [animal J7 right hemisphere]; Uninjected hemisphere [animal UW8]; Supp. Table 1). Scale bar = 500 um. d – i, Fluorescence labeling in striatal tissue (same animals as a – c) of tdTomato (red), the astrocyte marker GFAP (white), the microglial marker Iba1 (green), and the nuclear marker DAPI (blue). White boxes show the regions of interest (ROIs) drawn for analysis of mean fluorescence intensity. The gray lines indicate where in the low magnification (d – f, scale = 100 μm) image each high magnification image (g – I, scale = 100 μm) is found. White arrows = astrocyte. White arrowheads = microglia. j and k, mean fluorescence intensity of (j) GFAP and (k) Iba1 expression. Graphs show group median and interquartile range. No significant differences were found between groups. Individual channels were adjusted for brightness as needed (Supp. Table 4). DAPI, 4′,6-diamidino-2-phenylindole; GFAP, glial fibrillary acidic protein; Iba1, ionized calcium binding adaptor molecule 1; NC, nanocapsule.

To characterize the discrete cellularity observed with HE, coronal brain sections were immunolabeled against the astrocyte marker GFAP and the microglial marker ionized calcium binding adaptor molecule 1 (Iba1). In all hemispheres, GFAP+ and Iba1+ cells were present to varying degrees (Fig. 5d-i). In the striatum of uninjected hemispheres, GFAP+ cells were minimal, reflecting resident astroglia. Scattered Iba1+ cells indicated the presence of resting microglia typified by highly ramified small, circular cell bodies extending multiple thin and branching processes. In all injected hemispheres, astrocytes and microglia appeared mildly to moderately more abundant than in the uninjected hemispheres. Iba1 immunolabeling was increased in small areas similar to the regions of increased cellularity observed in HE, following the needle track. The Iba1+ microglia in these foci displayed a more ameboid, activated phenotype, appearing larger and with fewer extended processes. Despite the mild increases in visible GFAP immunolabeling and focal areas of increased Iba1immunolableing in the injected hemispheres, quantification of mean fluorescence intensity in striatum of these animals did not show statistically significant differences between treatment groups (GFAP: Ai14 sgRNA 166.9 ± 35.4, non-targeting sgRNA 154.3 ± 52.8, uninjected 95.8 ± 19.3; *H* = 5.422, *df* = 2, *p* = 0.0714; Iba1: Ai14 sgRNA 182.4 ± 23.8, non-targeting sgRNA 174.2 ± 47.0, uninjected 159.1 ± 24.9, *H* = 0.8, *df* = 2, *p* = 0.7214; Fig. 5j, k). The triple-immunolabeling also provided further confirmation of the preferential targeting of the NCs to neurons, as there was very little co-localization of tdTomato with GFAP or Iba1.

## DISCUSSION

Our results demonstrate successful *in vivo* genome editing in the brain following delivery of CRISPR RNP by our uniquely engineered biodegradable RNP NCs. NCs preferentially targeted neurons, with minimal editing in glial cells. Neuronal DNA editing was efficiently produced by the NC-delivered RNP, with about 26% of neurons in the target area expressing tdTomato. It is important to note that expression of the tdTomato protein in the Ai14 mouse model significantly underreports the actual genome editing efficiency. The Ai14 sgRNA has three target sites in the stop cassette and can produce edits such as small indels or a deletion of only a single stop sequence repeat, neither of which activate tdTomato expression. It has been estimated that only 34% to 40% of edited cells are expected to produce tdTomato ^29^, indicating that greater than 60% of neurons are likely edited in the present study.

The genome-edited cells were largely DARPP32+, post-mitotic, medium spiny neurons, a gamma-aminobutyric acid (GABA)-ergic neuron population that comprises 95% of the neurons in the striatum ^34^. Co-labeling of tdTomato with ChAT, parvalbumin, and calretinin confirmed the capability of NCs to produce editing across multiple neuronal phenotypes. The low incidence of genome editing in cholinergic, parvalbumin, and calretinin neurons reflects the low number of these interneuron subtypes in the rodent striatum, estimated to be 0.5-2% ^35,36^, 0.7% ^36^, and 0.5% ^36^, respectively.

These results and previously demonstrated NC-induced genome editing in HEK 293 cell culture, murine retinal pigmented epithelium (RPE), and murine muscle cells ^32^ showcase the utility of this CRISPR RNP delivery method across *in vitro* and *in vivo* applications. These NCs are particularly desirable for intracerebral delivery of RNP to neurons for several reasons. First, the small size of NCs permits them to move through the brain parenchyma and deliver cargo to murine striatal neuron s^37,38^. The significant increase in total edited brain area in the animals that received an injection volume of 1.5 µl compared to the animals that received 1 µl in this study is consistent with increased volume contributing to the spread of the NCs in brain tissue. This finding suggests that the injected volume can be adjusted as needed for local administration aiming to target specific brain structures and minimize concerns of editing in non-targeted brain areas, especially if combined with real time-intraoperative MRI targeting ^39^. A larger injection volume could be paired with techniques such as convection enhanced delivery ^40^ to increase NC distribution to generate a greater edited area. Second, the NC surface is highly PEGylated, which efficiently reduces surface charge and hydrophobicity, enabling fast diffusion within the brain extracellular matrix ^37,41^. Lastly, the NC surface can be easily modified with targeting ligands (e.g., RVG peptide) or CPP, which can potentially enhance the editing efficiency of the specific cell type being targeted. Interestingly, the addition of either CPP or RVG to the NC in the present study did not significantly alter the neuronal editing efficiency or size of the edited brain area. It is currently unclear why differences were not observed in the animal groups treated by different NC formulations. One potential explanation is that the targeting ligand type or molar ratio may need to be optimized to edit a greater number of neurons or additional neuron types. In previous work with these NCs, addition of the RPE targeting ligand all-trans retinoic acid (ATRA) produced significantly increased *in vivo* RPE editing relative to undecorated NCs ^32^, illustrating the utility of target ligand decoration on these NCs for specific cell types.

Several vehicles and methods for delivering CRISPR genome editors to the brain have been developed and evaluated. AAV vectors have been directly injected into numerous brain regions to edit neuronal proteins ^42-46^, including in rodent models of Huntington’s disease ^47,48^ and Alzheimer’s disease ^49^. Intracerebroventricular (ICV) delivery of AAV carrying CRISPR has been explored to maximize the area of genome editing in the brain due to distribution in the cerebrospinal fluid (CSF) ^16,17^. A recent study utilizing ICV delivery of AAVs showed knockdown of NeuN in multiple CNS regions ^16^. The degree of NeuN knockdown was noted as variable across the brain and spinal cord, probably due to limited penetration across the brain parenchyma from the cerebroventricular system. Furthermore, viral vector delivery of CRISPR gene editing components is subject to the limitations described in the introduction, including potential immune response. A gold nanoparticle-based, cationic polymer-coated nanocarrier, i.e., CRISPR-Gold has also been used to produce genome-editing in the brain following injection into the striatum or hippocampus ^30^. The CRISPR-Gold Cas9 delivery method is well suited for editing multiple cell types in the brain, particularly glial cells, as it seems to preferentially edit resident brain astroglia (approx. 60% of total edited cells) and microglia (approx. 25%) compared to neurons (approx. 15%) in the Ai9 mouse striatum. RNP delivery into the striatum or midbrain by nanocomplexes formed by an R7L10 peptide ^31^ has been shown to produce neuronal editing. The size of the edited brain area was not reported for this study, but R7L10 nanocomplexes are reported to have a larger size (approx. 100 nm) and positive charge (around 10 mV) which might limit their diffusion capability and, therefore, they may produce a small edited brain area.

NC administration and genome-editing were well tolerated by the animals in the current study and no appreciable immune response in the brain was identified. Injection of NC-encapsulated RNP carrying either Ai14 targeting or non-targeting sgRNA induced minimal increased cellularity in the brain parenchyma along the needle tract two weeks post NC delivery. Immunolabeling of astrocytes and microglia did not show a statistically significant difference in the mean fluorescence intensity of these glial markers between NC-injected and uninjected hemispheres. A mild to moderate increase in gliosis is typical following insertion of a needle into the brain and is observed following saline injection ^50^. Assessment of inflammation at an earlier timepoint, e.g. 7 days post-injection instead of 14 days, may have detected a greater immune response as astrogliosis resolves over time ^50^.

The RNP NCs produced robust *in vivo* neuronal editing, independently validated in separate experimental cohorts. The experiments produced consistent results while taking place at two different institutions (i.e., UW-Madison and The Jackson Laboratory) with separate surgical teams, imaging tools, and raters for ROI drawing and cell counting. The repeated demonstration of efficient editing of neurons in the brain, combined with previous results of *in vivo* editing in additional tissue types ^32^, showcases the effectiveness and versatility of the NCs as a CRISPR RNP delivery platform. The evaluation of the NCs at multiple facilities was made possible by the notable stability of these NCs, which will be critical for biomedical applications of this technology. In proper storage buffer (i.e., 20 mM HEPES-NaOH pH 7.5, 150 mM NaCl, 10 % glycerol), the NC was stable for 130 days at -80°C, indicating NC is suitable for long-term storage with good gene editing efficiency preserved.

Overall, these data provide important proof of principle of the efficacy and safety of the NCs in the mammalian brain. Experiments thus far have focused on the Ai14 mouse model, which is not disease relevant. An important next step for preclinical testing of these NCs will be to demonstrate editing efficiency and safety of a therapeutically relevant target in a nonhuman primate model. TdTomato-positive neurons were indistinguishable with respect to morphology from unedited cells, indicating healthy axons and dendrites and active, intact transcription and translational processes within the edited cells. While we did not perform functional studies on the edited mice or brain slices, we did not see any gross behavioral changes in the treated mice within the 2-3 week timescale of the experiments. These results are consistent with healthy function of the retinal and muscle tissue following injection of NCs in prior studies ^32^, and we expect any potential adverse effects in the edited neurons to be low and evaluated in future studies. In addition, scaling up the production of the NCs for administration to the larger animals will be needed. These future studies are warranted in order to progress toward clinical translation of the biodegradable NCs as a treatment for neurological diseases.

## METHODS

### Materials

Acrylic acid (AA), *N,N,N*′,*N*′-tetramethylethylenediamine (TEMED), 1-vinylimidazole (VI) and ammonium persulfate (APS), and tris(2-carboxyethyl)phosphine (TCEP)were purchased from Thermo Fisher Scientific. Acrylate-mPEG (Ac-mPEG, 2 kDa) and acrylate-PEG-maleimide (Ac-PEG-Mal, 2 kDa) were acquired from Biochempeg Scientific Inc. *N*-(3-aminopropyl)methacrylamide hydrochloride (APMA) and *N,N*′-bis(acryloyl)cystamine (BACA) were purchased from Sigma-Aldrich. Peptides, Cys-TAT (CYGRKKRRQRRR) and RVG-Cys (YTIWMPENPRPGTPCDIFTNSRGKRASNGC) were synthesized by Genscript. Nuclear localization signal (NLS)-tagged Streptococcus pyogenes Cas9 nuclease (sNLS-SpCas9-sNLS) was obtained from Aldevron. *In vitro* transcribed single guide RNAs (sgRNAs) were purchased from Integrated DNA Technologies, Inc., or Synthego. The sgRNAs used in this experiment include the Ai14 sgRNA (protospacer 5′ - AAGTAAAACCTCTACAAATG-3′) and a non-targeting sgRNA (Alt-R CRISPR-Cas9 Negative Control crRNA #1, Integrated DNA Technologies, Inc., USA). GFP-targeting sgRNAs (GFP protospacer: 5’-GCACGGGCAGCTTGCCGG-3’) were purchased from Synthego (i.e., sgRNA 1) and Integrated DNA Technologies (i.e., sgRNA 2).

### Synthesis of peptide conjugated Ac-PEG (Ac-PEG-CPP and Ac-PEG-RVG)

Ac-PEG-CPP and Ac-PEG-RVG were synthesized via a Michael addition reaction between Ac-PEG-Mal and corresponding peptides with cysteine terminals. Typically, 10 μmol of peptide was mixed with Ac-PEG-Mal (24 mg, 1.2 μmol) in DI water containing 5 mM TCEP, with the pH of the mixture adjusted to 7. The reaction was carried out under a nitrogen atmosphere at room temperature. After 12 h, the Ac-PEG-peptide was purified by dialysis against DI water for 48 h (MWCO 2kDa) and lyophilized to obtain the products in dry powder form. The ^1^H-NMR spectrum of Ac-PEG-CPP and Ac-PEG-RVG were shown in Supp.Figs. 4 and 5, respectively (D_2_O, 400 MHz).

### Preparation of Cas9-sgRNA ribonucleoprotein (RNP)

The sNLS-Cas9-sNLS protein was combined with sgRNA at a 1:1 molar ratio. The mixture was allowed to complex for 5 min on ice with gentle mixing. The as-prepared RNP was used freshly, without further purification.

### Preparation of NCs

NCs were prepared as previously reported with minor modifications ^32^. Prior to NC synthesis, pH = 8.5 sodium bicarbonate buffer (5 mM) was freshly prepared and degassed using the freeze–pump–thaw method for three cycles. Monomers, AA, APMA, VI and Ac-PEG were accurately weighed and dissolved in degassed sodium bicarbonate buffer (10 mg/ml). The crosslinker, BACA, was dissolved in DMSO (2 mg/ml). The free radical initiators, APS and TEMED were accurately weighed and dissolved in degassed sodium bicarbonate buffer (10 mg/ml). The Cas9 RNP was placed in a Schlenk flask and diluted to 0.12 mg/ml in sodium bicarbonate buffer in a nitrogen atmosphere. Monomer solutions were added into the above solution under vigorous stirring in the order of AA, APMA and VI at 5 min intervals. In each 5 min interval, the solution was degassed by vacuum pump for 3 min and refluxed with nitrogen. After another 5 min, the crosslinker, BACA, was added, followed by the addition of APS. The mixture was degassed for 5 min, and the polymerization reaction was immediately initiated by the addition of TEMED. The *in situ* free radical polymerization was carried out under a nitrogen atmosphere for 50 min. Thereafter, Ac-PEG was added. The solution was degassed by a vacuum pump, and the reaction was resumed for another 20 min to allow for NC surface PEGylation. The as-prepared NC was purified and concentrated by Ultrafiltration using Amicon ® Ultra centrifugal filters (MilliporeSigma, MWCO 100 kDa) and redispersed in NC storage buffer (20 mM HEPES-NaOH pH 7.5, 150 mM NaCl, 10 % glycerol). The molar ratio of AA/APMA/VI/BACA/Ac-PEG/RNP used for the optimal formulation was 927/927/244/231/64/1. The weight ratio of RNP/APS/TEMED was kept at 1/0.5/0.5, corresponding to a molar ratio of approximately 1/350/700. NC-CPP and NC-RVG were prepared following a similar protocol as described above with the molar ratio of AA/APMA/VI/BACA/Ac-mPEG/Ac-PEG-CPP(or RVG) at 927/927/244/231/32/32.

### Characterization

The sizes and zeta-potentials of NCs were studied by dynamic light scattering (ZetaSizer Nano ZS90). NCs were redispersed in DI water, and the pH was adjusted to 7.4 by 1 M NaOH, prior to DLS and zeta potential measurements. The NC concentrations for DLS and zeta potential were 0.1 mg/ml and 0.05 mg/ml, respectively. The morphologies of NCs were also characterized by transmission electron microscopy (TEM, FEI Tecnai 12, 120 keV).

### Cell culture and NC storage studies

GFP-expressing human embryonic kidney cells (i.e., GFP-HEK cells, GenTarget Inc.) were used as an RNP delivery cell model. GFP-HEK cells were cultured with DMEM medium (Gibco, USA) added with 10% (v/v) fetal bovine serum (FBS, Gibco, USA) and 1% (v/v) penicillin–streptomycin (Gibco, USA). Cells were cultured in an incubator (Thermo Fisher, USA) at 37°C with 5% carbon dioxide at 100% humidity.

For the storage test, NCs with GFP-targeting sgRNAs (GFP protospacer: 5’-GCACGGGCAGCTTGCCGG-3’) purchased from Synthego (i.e., sgRNA 1) and Integrated DNA Technologies (i.e., sgRNA 2) were prepared and redispersed in NC storage buffer with a RNP concentration of 20 μM. The NCs were then aliquoted and stored at different temperatures (i.e., 4°C, -20 and -80°C) in a storage buffer (20 mM HEPES-NaOH pH 7.5, 150 mM NaCl,10 % glycerol). The RNP NCs were flash-frozen in liquid nitrogen prior to storage at -20 °C or -80 °C. GFP-HEK cells were seeded at a density of 5,000 cells per well onto a 96-well plate 24 h prior to NC treatments. Cells were treated with NCs with a RNP dose of 150 ng/well (or an equivalent Cas9 protein dose of 125 ng/well). After 96 h, cells were detached from the well plates with 0.25% trypsin-EDTA, spun down and resuspended in 500 μl phosphate buffered saline (PBS). The editing efficiency was assayed using flow cytometry by quantifying the percentage of GFP-negative cells.

### Intracerebral Injections

Ai14 mice (The Jackson Laboratory (JAX), STOCK# 7914) were used to assay the gene editing efficiency in the brain (Fig. 2a). Experiments were conducted at both UW-Madison and JAX as part of the NIH Somatic Cell Genome Editing Consortium (SCGE) effort to show repeatability of findings. See Supp. Table 1 for details on subjects, assigned treatments, and tissue used for analysis.

All animal treatments and procedures were approved by either the University of Wisconsin–Madison or JAX Animal Care and Use Committee as appropriate. Mice were examined and determined to be in good health the day of injection. Mice were anaesthetized by either intraperitoneal injection of a ketamine (120 mg/kg), xylazine (10 mg/kg), and acepromazine (2 mg/kg) cocktail (UW-Madison) or isofluorane gas (inhalation to effect, typically 1-3%) (JAX). Stereotactic brain injections were performed using a Stoelting stereotaxic frame equipped with a Stoelting Quintessential Stereotax Injector (QSI). Solutions were intracerebrally delivered at a rate of 0.2 μl/minute. After the injection was completed, the needle remained in place for up to 5 minutes, then the surgical field was irrigated with sterile saline and the skin layers closed with surgical glue.

For the UW-Madison methods development animal cohort, the right and/or left striatum was targeted at coordinates of AP +0.74 mm, ML ±1.74 mm, DV -3.37 mm using a 10 μl Hamilton syringe and 32-gauge 1 inch Hamilton small hub RN needle. The solutions delivered were 1.5 μl of NC-No Ligand or NC-CPP with RNP carrying guide targeting either Ai14 or a non-targeting guide at 20 μM RNP suspended in PBS or 1 μl of storage buffer (Supp. Table 1). Brain hemispheres of mice injected with NCs with RNP containing Ai14 targeting guide in the striatal target area were imaged for neuron editing analysis (NC-CPP, n = 3 hemispheres; NC-No Ligand, n = 4 hemispheres). In addition, uninjected mouse brain hemispheres (n = 3) were also imaged to assess host reaction to the NCs (Supp. Table 1).

For JAX animal cohorts, the right striatum was targeted at coordinates AP +1.2 mm, ML+/- 1.6 mm, DV -3.37) (Fig. 2b; Supp. Table 1), using a similar syringe set up. The solutions delivered were 1 μl of NC-No Ligand (n = 6 hemispheres), NC-RVG (n = 6 hemispheres), or NC-CPP (n = 6 hemispheres) carrying guide targeting Ai14 at 20 μM RNP suspended in a storage buffer. The same volume of NCs carrying the non-targeting guide was injected into the contralateral hemisphere.

### Necropsy and tissue processing

For all animals, brain tissue was collected two weeks after intracerebral injection. At UW-Madison, mice were deeply anesthetized with a combination of ketamine (120 mg/kg) and xylazine (10 mg/kg) and transcardially perfused with heparinized saline. Brains were retrieved, post-fixed for 24 hours in 4% PFA, and cryoprotected in graded sucrose. At JAX, mice were euthanized via CO2 asphyxiation, followed immediately by transcardiac perfusion with heparinized PBS. Brains were collected, post-fixed for 48 hours in 4% PFA, and cryoprotected in 30% sucrose/PBS at 4°C for 48 hours. All brains were cut frozen in 40 μm coronal sections on a sliding microtome and stored at -20°C in cryoprotectant solution until staining. For UW-Madison animals, while cutting, the left hemisphere of each coronal slice was identified by making two small holes in the cerebral cortex. The cryoprotectant solution at UW-Madison consisted of 1000 ml 1X PBS (pH 7.4), 600 g sucrose, 600 ml ethylene glycol and at JAX of 350 ml 0.1 M PB solution (pH 7.35), 150 ml ethylene glycol, 100 μg sodium azide, 150 g sucrose.

### Anatomical Evaluation

Serial coronal brain sections spaced 240 μm apart from three JAX injected NC-No Ligand group animals and one animal with an uninjected brain hemisphere were stained for HE and blindly evaluated by a board-certified veterinary pathologist. The sections were assessed for histological changes such as presence and severity of inflammation or atrophy. Striatal sections from one animal with an uninjected brain hemisphere were stained for Nissl to collect an image illustrating normal murine striatal anatomy (Fig. 2c).

### Immunohistochemistry

All immunolabeling was performed using one sixth serial brain sections spaced 240 μm apart from each animal to sample the entire rostrocaudal span of the striatum. After three 10-minute washes in Tris buffered saline (TBS) plus 0.05% TritonX-100, background staining was blocked with a 2 hour incubation in a (TBS) solution containing 5% normal serum, 2% bovine serum albumin, and 0.05% Triton X-100. The slices were incubated with primary antibody (Supp. Table 2) for 24 hours at room temperature, washed 3 times in dilution media, and then incubated for 2 hours at room temperature with secondary antibody (Supp. Table 3). After three 10-minute washes in PBS, the tissue was counterstained with DAPI, mounted onto slides, allowed to dry, and coverslipped with Fluor Gel. Immunostaining of tissue sections from animals in different treatment groups was performed in parallel and included negative and positive controls. Positive controls for tdTomato consisted of tissue from an Ai9 mouse (JAX 7909) crossed with an Myf-Cre expressing mouse (JAX, 7893, tissue provided by Murray Lab). Negative controls were performed by omitting primary antibodies.

### Image acquisition

Image acquisition performed at UW-Madison utilized a Nikon A1R confocal microscope with 405, 488, 561, and 640 wavelength lasers using NIS Elements version 5.20.02. Detectors for the 488 and 561 lasers are high sensitivity GaAsP PMTs, while the 405 and 640 lasers use HS PMTs. Whole coronal brain slice images were acquired using the 4x objective (Plan Apo, N.A. = 0.2, Nikon) and using XY stitching with 30-35% overlap. Images used for analysis of neuronal editing efficiency/edited area (methods development cohort), types of edited neurons, or glial response were acquired at UW-Madison using the 20x objective (Plan Apo VC, N.A. = 0.75, Nikon) with XY stitching with 30-35% overlap in either a single focal plane (glial response) or in 3 focal planes each 5 µm apart covering a total of 10 µm (neuronal editing efficiency/editing area and types of edited neurons). High magnification images to show details of colabeling were acquired at 40x (Plan Fluor, N.A. = 0.75, Nikon) with multiple focal planes. The size of each frame was 1024 × 1024 pixels, and the intensity of the signal in each pixel was recorded at 12-bits for each channel. Images taken in multiple focal planes were processed as maximum intensity projections for figures and analysis.

Image acquisition performed at JAX used a Leica DMi8 widefield microscope (to evaluate genome-edited edited area size) and a Leica Sp8-AOBS confocal microscope equipped with 405, 458, 488, 514, 561, 594, and 633 nm wavelength lasers and the LASX software (to evaluate neuronal editing efficiency). The Leica Sp8-AOBS confocal microscope is equipped with UV/DAPI (A), FITC/AF-488/GFP (I3), Tritc/Rhod/DsRed (N2.1) filters. PMT detectors are fed by an acousto-optical beam splitter (AOBS) and a spectral detector (prism). Images were acquired using the 20x objective (NA0.75 GLYC WD = 0.66 mm CORR) with XY stitching with 10% overlap in 3 focal planes each 5 µm apart covering a total of 10 µm. The size of each frame was 1024-2079 × 1024-3947 and the intensity in each pixel was recorded at 16-bits for each channel.

During the preparation of images for figures for the manuscript, any adjustments to image brightness, such as adjusting of LUTs of immunofluorescent images, were applied to the entire image. Images with adjustments to individual channel LUTs have this noted in figure legends with detailed information in Supp. Table 4.

### Image analysis

All analysis of neuronal editing efficiency and genome-edited area size, for both the UW-Madison and JAX injected animals, was performed using tdTomato/NeuN/GFAP triple-immunolabeled tissue counterstained with DAPI (Supp. Table 1). ROIs were drawn in maximum intensity projection images around areas of tdTomato signal in which a group of at least 5 cells were tdTomato+ and within 135 μm of each other. Neuronal editing efficiency was calculated for the three largest ROIs for each animal, as the percentage of NeuN+ cells that were also tdTomato+.

In the UW-Madison injected animals, ROIs were drawn in NIS Elements version 5.30.02 using the Draw Polygonal ROI function in the red (tdTomato) channel of the maximum intensity projection image, and the size (area in μm^2^) was exported for each ROI. Using NIS Elements, the total number of NeuN+ cells inside the ROI was calculated using the Binary Function followed by manual correction. A threshold was defined in the NeuN channel with the lower range set to exclude background, 3x smooth, 6x clean, 1x separate, and size selection > 5 μm. The automated count was then manually edited based on NeuN immunolabeling to split groups of cells counted as a single cell and to exclude NeuN+ cells with less than 50% of the cell soma inside the ROI. Genome-edited neurons were defined as NeuN+ and tdTomato+ and were manually selected in NIS Elements and counted.

In JAX injected animals, polygonal ROIs were drawn in FIJI following duplication of the tdTomato channel and conversion to a binary, and the size (area in μm^2^) was exported for each ROI and recorded (Fig. 2). To count the total number of neurons, the NeuN channel was duplicated and converted to a binary. Images were then processed using the StarDist2D plugin with fluorescent default settings. The threshold of the resulting Label Image (V) was adjusted so that all cells were visible, and the image was processed using the watershed tool to separate touching objects. The ROI was then copied from the tdTomato image and NeuN+ cells counted using the Analyze Particle function with size 40-infinity and circularity 0-1 (Fig. 2). Automated NeuN counts using this method significantly correlated with manual counts performed in a subset of images (*ρ* = 0.897, *p* <0.0001; Supp. Table 5). Genome-edited NeuN+/tdTomato+ neurons within each ROI were manually selected and counted using the multipoint tool in FIJI (Fig. 2).

Assessing the types of neurons that were genome-edited was performed with tdTomato/DARPP32/ChAT or tdTomato/Parvalbumin/Calretinin triple-immunolabeled tissue counterstained with DAPI (n=3 per NC treatment group; Supp. Table 1). Neuron counts were performed using the Taxonomy function in NIS Elements version 5.30.02 in 800 μm x 1200 μm ROIs in 2-4 coronal tissue sections per animal.

For analysis of glial response, imaging was performed on tdTomato/GFAP/Iba1 triple-immunolabeled tissue counterstained with DAPI (Supp. Table 1). Mean fluorescence intensity of 2 coronal tissue sections were evaluated per animal to compare uninjected, NC-No Ligand non-targeting sgRNA, and NC-No Ligand Ai14 sgRNA groups (n=3 hemispheres per group). In each section, data was averaged from two nonoverlapping ROIs with ROI size of 500 µm x 500 µm. ROIs were placed around the areas with the highest GFAP or Iba1 immunolabeling in the target area.

### Statistics

All statistical analyses were performed in GraphPad Prism v9. Comparisons between groups were made using the Kruskal-Wallis test (test statistic = *H*) with Dunn’s test for multiple comparisons for tests involving three or more groups (JAX animals editing efficiency treatment group comparison, JAX animals edited area treatment group comparison, edited neuron types treatment group comparison, Iba1 and GFAP mean fluorescence intensity). For multiple comparisons tests involving two groups (UW-Madison animals editing efficiency treatment group comparison and UW-Madison animals edited area treatment group comparison) the Mann-Whitney test (test statistic = *U*) was performed and reported. *In vivo* animal averages are presented in the text as median ± interquartile to be consistent with the nonparametric statistical tests performed due to multiple datasets exhibiting non-normal distributions. Spearman correlation was used to test the relationship between automated vs. manual NeuN counts. All *p* values reported are two tailed, and a *p* value < 0.05 is considered statistically significant. Effect sizes are given for statistically significant *p* values. Effect size for Spearman correlation is reported as the test statistic *ρ*. Effect sizes for Mann-Whitney are reported as *r* = (*Z*/(sqrt(*n*))) where *r* is the effect size, *Z* is the standardized test statistic, and n is the total number of observations.

## ACKNOWLEDGEMENTS

We are grateful for the technical support provided by Kat Jones and Pankaj Dubey. We thank the highly trained animal care technicians, surgeons, and colony managers at JAX for their hard work and dedication with special thanks to Randy Walls, Amy Leighton, and Jonathan Newell. Additional thanks to Lawrence Bechtel, Seth Hannigan and Jeff Duryea, Jr. at the JAX Small Animal Testing Center, for their talents in collecting the data associated with this and similar SCGE consortium validation projects. This work was supported by the NIH Common Fund and National Institutes of Health Office of the Director U54 OD026635 (S.A.M. and C.M.L), the WIMR optical imaging core (1S10OD025040-01), NIH 1-UG3-NS-111688-01 (S.R., K.S., J.L., M.E.E., S.G), and NIH UH3 grant # 4-UH3-NS111688 (S.R., K.S., J.L., M.E.E., S.G).

## DATA SHARING PLAN

The datasets generated during and/or analyzed during the current study are available from the corresponding authors on reasonable request.

## REFERENCES

1 Sharma, G., Sharma, A. R., Bhattacharya, M., Lee, S.-S. & Chakraborty, C. CRISPR-Cas9: A Preclinical and Clinical Perspective for the Treatment of Human Diseases. Molecular Therapy 29, 571–586, doi:10.1016/j.ymthe.2020.09.028 (2021).

2 Saha, K. et al. The NIH Somatic Cell Genome Editing program. Nature 592, 195–204, doi:10.1038/s41586-021-03191-1 (2021).

3 Stadtmauer, E. A. et al. CRISPR-engineered T cells in patients with refractory cancer. Science 367, eaba7365, doi:doi:10.1126/science.aba7365 (2020).

4 Gillmore, J. D. et al. CRISPR-Cas9 In Vivo Gene Editing for Transthyretin Amyloidosis. The New England journal of medicine 385, 493–502, doi:10.1056/NEJMoa2107454 (2021).

5 Maeder, M. L. et al. Development of a gene-editing approach to restore vision loss in Leber congenital amaurosis type 10. Nature medicine 25, 229–233, doi:10.1038/s41591-018-0327-9 (2019).

6 Doudna, J. A. Genomic engineering and the future of medicine. Jama 313, 791–792, doi:10.1001/jama.2015.287 (2015).

7 Heidenreich, M. & Zhang, F. Applications of CRISPR-Cas systems in neuroscience. Nature reviews. Neuroscience 17, 36–44, doi:10.1038/nrn.2015.2 (2016).

8 Bourasset, F., Auvity, S., Thorne, R. G. & Scherrmann, J. M. Brain Distribution of Drugs: Brain Morphology, Delivery Routes, and Species Differences. Handb Exp Pharmacol, doi:10.1007/164_2020_402 (2021).

9 Pardridge, W. M. Blood-Brain Barrier and Delivery of Protein and Gene Therapeutics to Brain. Frontiers in aging neuroscience 11, doi:10.3389/fnagi.2019.00373 (2020).

10 Kimura, S. & Harashima, H. Current Status and Challenges Associated with CNS-Targeted Gene Delivery across the BBB. Pharmaceutics 12, doi:10.3390/pharmaceutics12121216 (2020).

11 Peviani, M. et al. Biodegradable polymeric nanoparticles administered in the cerebrospinal fluid: Brain biodistribution, preferential internalization in microglia and implications for cell-selective drug release. Biomaterials 209, 25–40, doi:10.1016/j.biomaterials.2019.04.012 (2019).

12 MacKay, J. A., Deen, D. F. & Szoka, F. C., Jr. Distribution in brain of liposomes after convection enhanced delivery; modulation by particle charge, particle diameter, and presence of steric coating. Brain research 1035, 139–153, doi:10.1016/j.brainres.2004.12.007 (2005).

13 Wolak, D. J. & Thorne, R. G. Diffusion of Macromolecules in the Brain: Implications for Drug Delivery. Molecular Pharmaceutics 10, 1492–1504, doi:10.1021/mp300495e (2013).

14 Colasante, G. et al. In vivo CRISPRa decreases seizures and rescues cognitive deficits in a rodent model of epilepsy. Brain 143, 891–905, doi:10.1093/brain/awaa045 (2020).

15 Zuckermann, M. et al. Somatic CRISPR/Cas9-mediated tumour suppressor disruption enables versatile brain tumour modelling. Nature communications 6, 7391, doi:10.1038/ncomms8391 (2015).

16 Hana, S. et al. Highly efficient neuronal gene knockout in vivo by CRISPR-Cas9 via neonatal intracerebroventricular injection of AAV in mice. Gene Therapy 28, 646–658, doi:10.1038/s41434-021-00224-2 (2021).

17 Torregrosa, T. et al. Use of CRISPR/Cas9-mediated disruption of CNS cell type genes to profile transduction of AAV by neonatal intracerebroventricular delivery in mice. Gene Therapy 28, 456–468, doi:10.1038/s41434-021-00223-3 (2021).

18 Conniot, J., Talebian, S., Simões, S., Ferreira, L. & Conde, J. Revisiting gene delivery to the brain: silencing and editing. Biomater Sci 9, 1065–1087, doi:10.1039/d0bm01278e (2021).

19 Duan, L. et al. Nanoparticle Delivery of CRISPR/Cas9 for Genome Editing. Frontiers in Genetics 12, doi:10.3389/fgene.2021.673286 (2021).

20 Bulcha, J. T., Wang, Y., Ma, H., Tai, P. W. L. & Gao, G. Viral vector platforms within the gene therapy landscape. Signal transduction and targeted therapy 6, 53, doi:10.1038/s41392-021-00487-6 (2021).

21 Hanlon, K. S. et al. High levels of AAV vector integration into CRISPR-induced DNA breaks. Nature communications 10, 4439, doi:10.1038/s41467-019-12449-2 (2019).

22 Li, A. et al. A Self-Deleting AAV-CRISPR System for In Vivo Genome Editing. Mol Ther Methods Clin Dev 12, 111–122, doi:10.1016/j.omtm.2018.11.009 (2019).

23 Li, L., Hu, S. & Chen, X. Non-viral delivery systems for CRISPR/Cas9-based genome editing: Challenges and opportunities. Biomaterials 171, 207–218, doi:10.1016/j.biomaterials.2018.04.031 (2018).

24 Chew, W. L. et al. A multifunctional AAV–CRISPR–Cas9 and its host response. Nature Methods 13, 868–874, doi:10.1038/nmeth.3993 (2016).

25 Weber, T. Anti-AAV Antibodies in AAV Gene Therapy: Current Challenges and Possible Solutions. Frontiers in immunology 12, doi:10.3389/fimmu.2021.658399 (2021).

26 Kim, S., Kim, D., Cho, S. W., Kim, J. & Kim, J. S. Highly efficient RNA-guided genome editing in human cells via delivery of purified Cas9 ribonucleoproteins. Genome research 24, 1012–1019, doi:10.1101/gr.171322.113 (2014).

27 Liang, X. et al. Rapid and highly efficient mammalian cell engineering via Cas9 protein transfection. Journal of biotechnology 208, 44–53, doi:https://doi.org/10.1016/j.jbiotec.2015.04.024 (2015).

28 Foss, D. V. & Wilson, R. C. Emerging Strategies for Genome Editing in the Brain. Trends in Molecular Medicine 24, 822–824, doi:10.1016/j.molmed.2018.07.008 (2018).

29 Staahl, B. T. et al. Efficient genome editing in the mouse brain by local delivery of engineered Cas9 ribonucleoprotein complexes. Nat Biotechnol 35, 431–434, doi:10.1038/nbt.3806 (2017).

30 Lee, B. et al. Nanoparticle delivery of CRISPR into the brain rescues a mouse model of fragile X syndrome from exaggerated repetitive behaviours. Nature biomedical engineering 2, 497–507, doi:10.1038/s41551-018-0252-8 (2018).

31 Park, H. et al. In vivo neuronal gene editing via CRISPR-Cas9 amphiphilic nanocomplexes alleviates deficits in mouse models of Alzheimer’s disease. Nature neuroscience 22, 524–528, doi:10.1038/s41593-019-0352-0 (2019).

32 Chen, G. et al. A biodegradable nanocapsule delivers a Cas9 ribonucleoprotein complex for in vivo genome editing. Nat Nanotechnol 14, 974–980, doi:10.1038/s41565-019-0539-2 (2019).

33 Madisen, L. et al. A robust and high-throughput Cre reporting and characterization system for the whole mouse brain. Nature neuroscience 13, 133–140, doi:10.1038/nn.2467 (2010).

34 Arlotta, P., Molyneaux, B. J., Jabaudon, D., Yoshida, Y. & Macklis, J. D. <em>Ctip2</em> Controls the Differentiation of Medium Spiny Neurons and the Establishment of the Cellular Architecture of the Striatum. The Journal of Neuroscience 28, 622–632, doi:10.1523/jneurosci.2986-07.2008 (2008).

35 Lemos, J. C., Shin, J. H. & Alvarez, V. A. Striatal Cholinergic Interneurons Are a Novel Target of Corticotropin Releasing Factor. The Journal of Neuroscience 39, 5647–5661, doi:10.1523/jneurosci.0479-19.2019 (2019).

36 Tepper, J., Tecuapetla, F., Koos, T. & Ibanez-Sandoval, O. Heterogeneity and Diversity of Striatal GABAergic Interneurons. Frontiers in Neuroanatomy 4, doi:10.3389/fnana.2010.00150 (2010).

37 Thorne, R. G. & Nicholson, C. In vivo diffusion analysis with quantum dots and dextrans predicts the width of brain extracellular space. Proceedings of the National Academy of Sciences 103, 5567–5572 (2006).

38 Nance, E. et al. Brain-penetrating nanoparticles improve paclitaxel efficacy in malignant glioma following local administration. ACS nano 8, 10655–10664 (2014).

39 Vermilyea, S. C. et al. Real-Time Intraoperative MRI Intracerebral Delivery of Induced Pluripotent Stem Cell-Derived Neurons. Cell transplantation 26, 613–624, doi:10.3727/096368916X692979 (2017).

40 Mehta, A. M., Sonabend, A. M. & Bruce, J. N. Convection-Enhanced Delivery. Neurotherapeutics : the journal of the American Society for Experimental NeuroTherapeutics 14, 358–371, doi:10.1007/s13311-017-0520-4 (2017).

41 Nance, E. A. et al. A dense poly(ethylene glycol) coating improves penetration of large polymeric nanoparticles within brain tissue. Sci Transl Med 4, 149ra119, doi:10.1126/scitranslmed.3003594 (2012).

42 Swiech, L. et al. In vivo interrogation of gene function in the mammalian brain using CRISPR-Cas9. Nat Biotechnol 33, 102–106, doi:10.1038/nbt.3055 (2015).

43 Platt, Randall J. et al. CRISPR-Cas9 Knockin Mice for Genome Editing and Cancer Modeling. Cell 159, 440–455, doi:https://doi.org/10.1016/j.cell.2014.09.014 (2014).

44 Yamaguchi, H., Hopf, F. W., Li, S. B. & de Lecea, L. In vivo cell type-specific CRISPR knockdown of dopamine beta hydroxylase reduces locus coeruleus evoked wakefulness. Nature communications 9, 5211, doi:10.1038/s41467-018-07566-3 (2018).

45 Nishiyama, J., Mikuni, T. & Yasuda, R. Virus-Mediated Genome Editing via Homology-Directed Repair in Mitotic and Postmitotic Cells in Mammalian Brain. Neuron 96, 755-768.e755, doi:10.1016/j.neuron.2017.10.004 (2017).

46 Sun, H. et al. Development of a CRISPR-SaCas9 system for projection- and function-specific gene editing in the rat brain. Science Advances 6, eaay6687, doi:doi:96cl:38010.1126/sciadv.aay6687 (2020).

47 Jing, L. et al. Accumulation of Endogenous Mutant Huntingtin in Astrocytes Exacerbates Neuropathology of Huntington Disease in Mice. Molecular neurobiology 58, 5112–5126, doi:10.1007/s12035-021-02451-5 (2021).

48 Yang, S. et al. CRISPR/Cas9-mediated gene editing ameliorates neurotoxicity in mouse model of Huntington’s disease. The Journal of clinical investigation 127, 2719–2724, doi:10.1172/JCI92087 (2017).

49 Sun, J. et al. CRISPR/Cas9 editing of APP C-terminus attenuates β-cleavage and promotes α-cleavage. Nature communications 10, 53, doi:10.1038/s41467-018-07971-8 (2019).

50 Lai, T. T. et al. Temporal Evolution of Inflammation and Neurodegeneration With Alpha-Synuclein Propagation in Parkinson’s Disease Mouse Model. Frontiers in Integrative Neuroscience 15, doi:10.3389/fnint.2021.715190 (2021).

