## Supplementary Figures 1-5, Supplementary Tables 1-5 for "Efficient in vivo neuronal genome editing in the mouse brain using nanocapsules containing CRISPR-Cas9 ribonucleoproteins"

Supplementary Materials

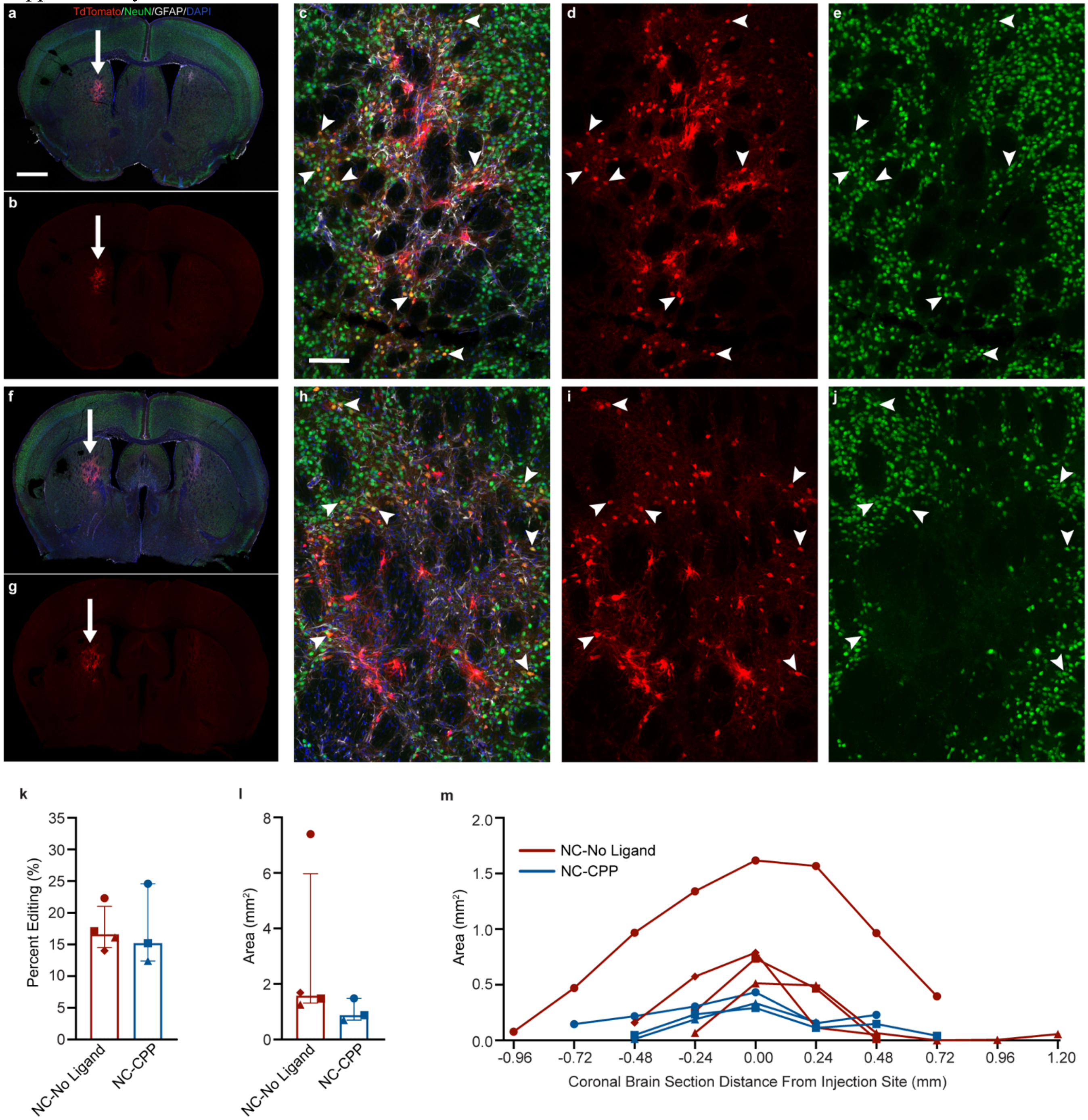


**Supp. Figure 1. Efficient *in vivo* neuronal genome editing is observed following injection of CRISPR RNP NCs in the methods development animal cohort (UW-Madison).** a, b, f, g, Coronal mouse brain sections showing neuronal genome-editing (white arrows) in (a,b) a representative NC-CPP treated mouse (animal UW1; Supp. Table 1) and (f,g) a NC-No Ligand treated mouse (animal UW2). c-e and h-j, Genome-edited neurons (yellow; white arrowheads) in the striatum (same animals as a, b, f, g). Scale bar = 100 µm. k, Percentage of neurons in the edited area expressing tdTomato in each NC treatment group. l, Sum of edited area size (region of interest area) across all coronal slices in each NC treatment group. k and l, Graphs show median and interquartile range. Differences between groups were not statistically significant. m, Line graph of edited area size for each individual animal at given distances rostral and caudal to the injection site. k – m, Each animal is shown with a unique color and symbol (Supp. Table 1). Photomicrographs show maximum intensity projection of three focal planes covering 10 µm. Individual channels were adjusted for brightness as needed (Supp. Table 4). CPP, cell penetrating peptide; DAPI, 4′ ,6-diamidino-2-phenylindole; GFAP, glial fibrillary acidic protein; NC, nanocapsule; NeuN, neuronal nuclear protein.

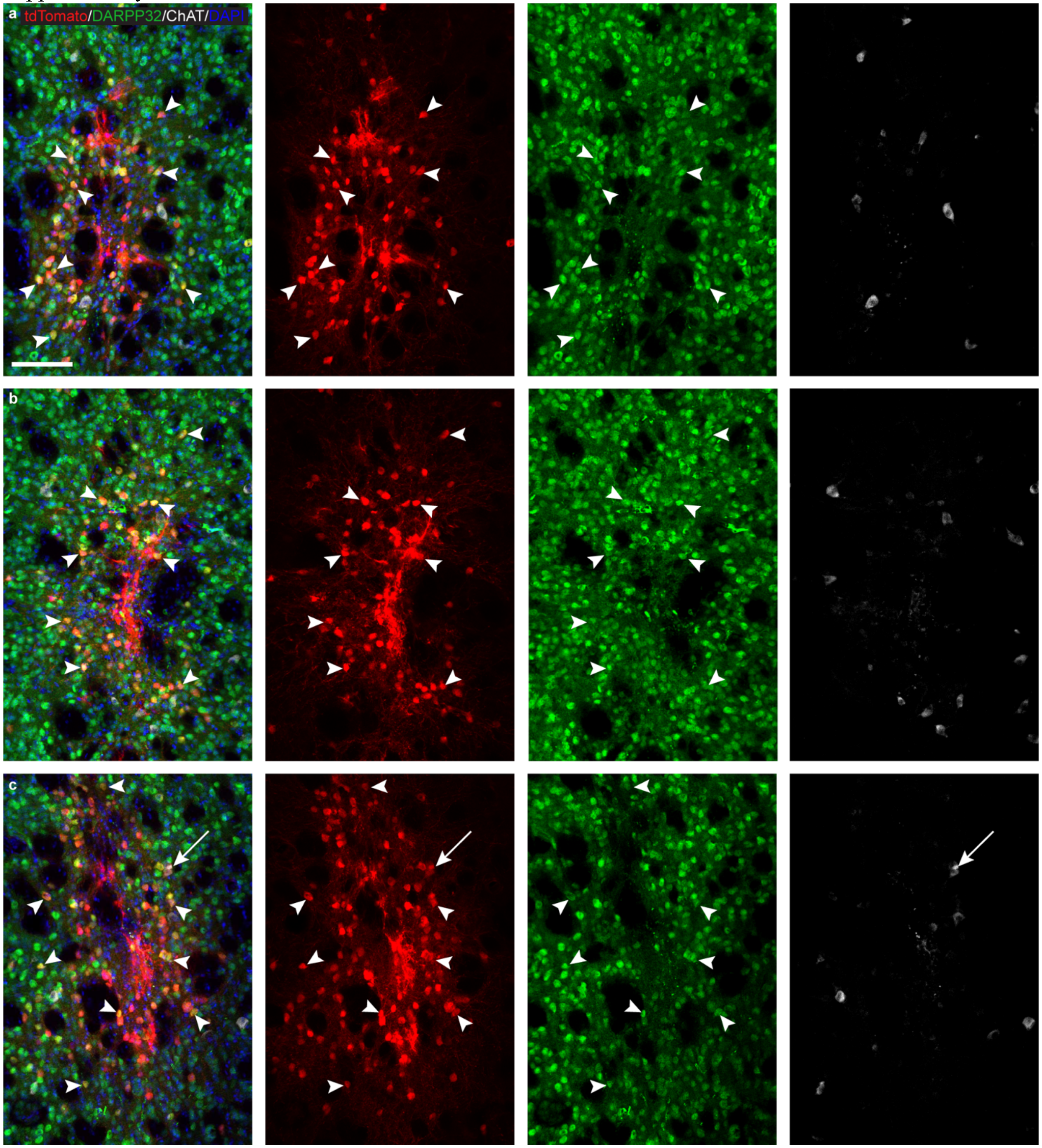


**Supp. Figure 2. Genome-edited neurons in the Ai14 mouse striatum following CRISPR RNP NC delivery were primarily medium spiny neurons and occasionally cholinergic interneurons.** The tdTomato+ (red) genome-edited neurons in the mouse striatum largely co-label with the medium spiny neuron marker DARPP32 (green; white arrowheads). A small number of tdTomato+ neurons also label for the cholinergic neuron marker choline acetyl transferase (ChAT; white; white arrows). This pattern was similar across NC formulations (a, NC-No Ligand animal J9; b, NC-CPP animal J17; NC-RVG animal J5; Supp. Table 1). Scale = 100 µm. Photomicrographs show maximum intensity projection of three focal planes covering 10µm. Individual channels were adjusted for brightness as needed (Supp. Table 4). CPP, cell penetrating peptide; DAPI, 4′,6-diamidino-2-phenylindole; DARPP32, dopamine- and cyclic-AMP-regulated phosphoprotein of molecular weight 32 kDa; NC, nanocapsule; RVG, rabies virus glycoprotein; RNP, ribonucleoprotein.

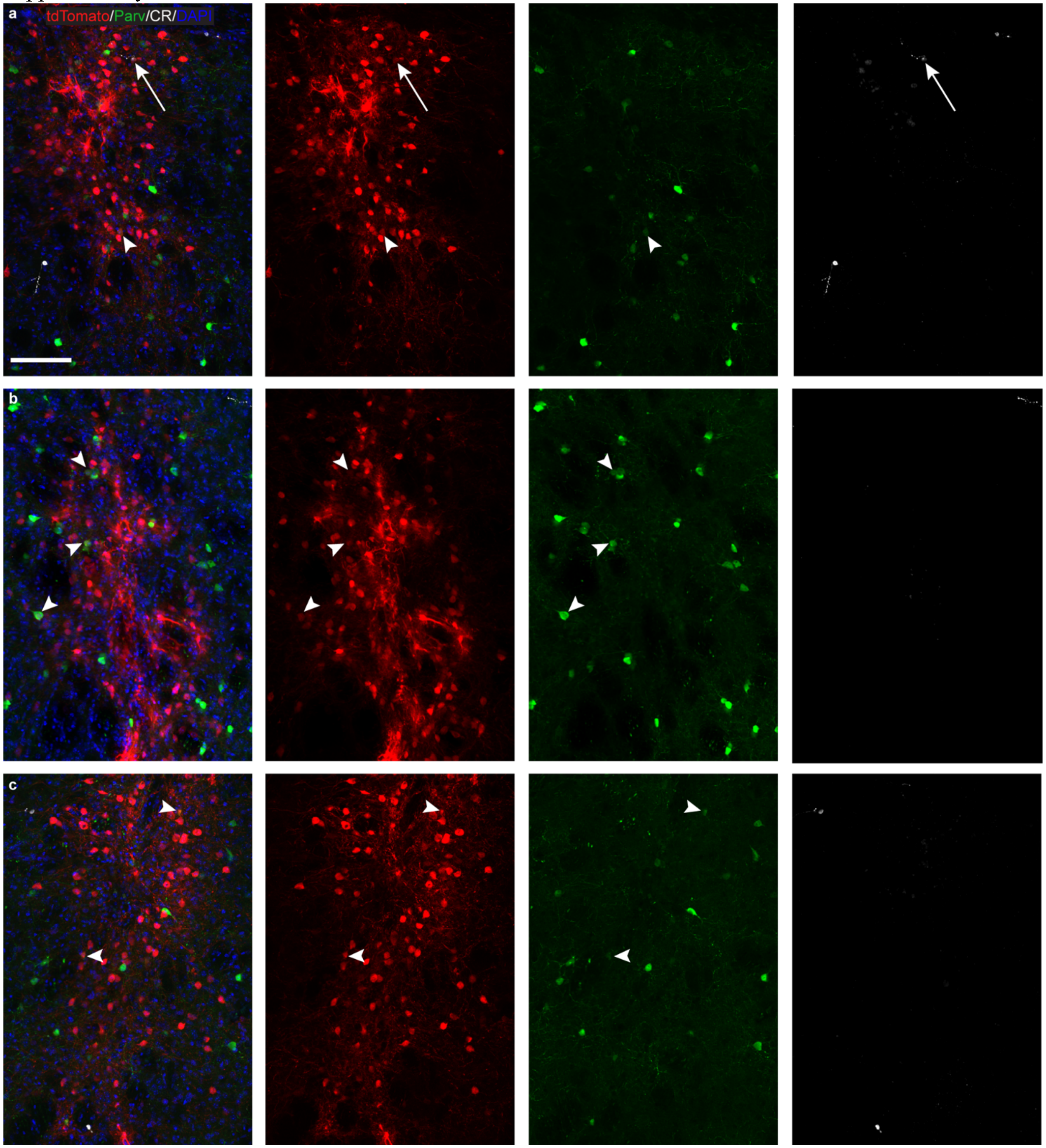


**Supp. Figure 3. Genome-edited neurons in the Ai14 mouse striatum following CRISPR RNP NC delivery were occasionally parvalbumin and calretinin neurons.** The tdTomato+ (red) genome-edited neurons in the mouse striatum occasionally co-label for parvalbumin (Parv, green; white arrowheads) or calretinin (CR, white; white arrows). This was similar across NC formulations (a, NC-No Ligand animal J8; b, NC-CPP animal J13; NC-RVG animal J4; Supp. Table 1). Scale = 100 µm. Photomicrographs show maximum intensity projection of three focal planes covering 10µm. Individual channels were adjusted for brightness as needed (Supp. Table 4). CPP, cell penetrating peptide; DAPI, 4′,6-diamidino-2-phenylindole; NC, nanocapsule; RVG, rabies virus glycoprotein; RNP, ribonucleoprotein.

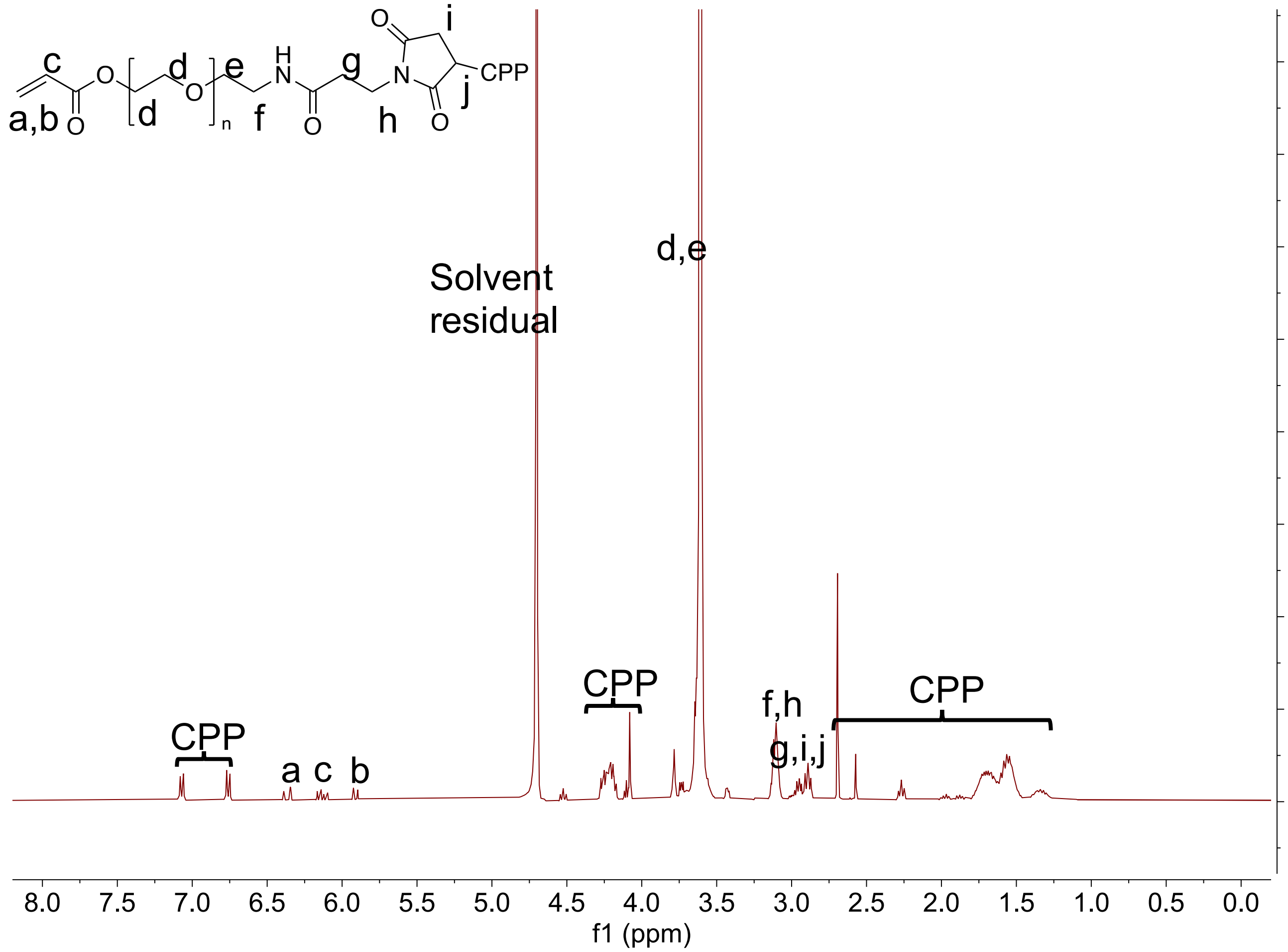


**Supp. Figure 4.** $^{1}$H NMR spectrum of Ac-PEG-CPP.

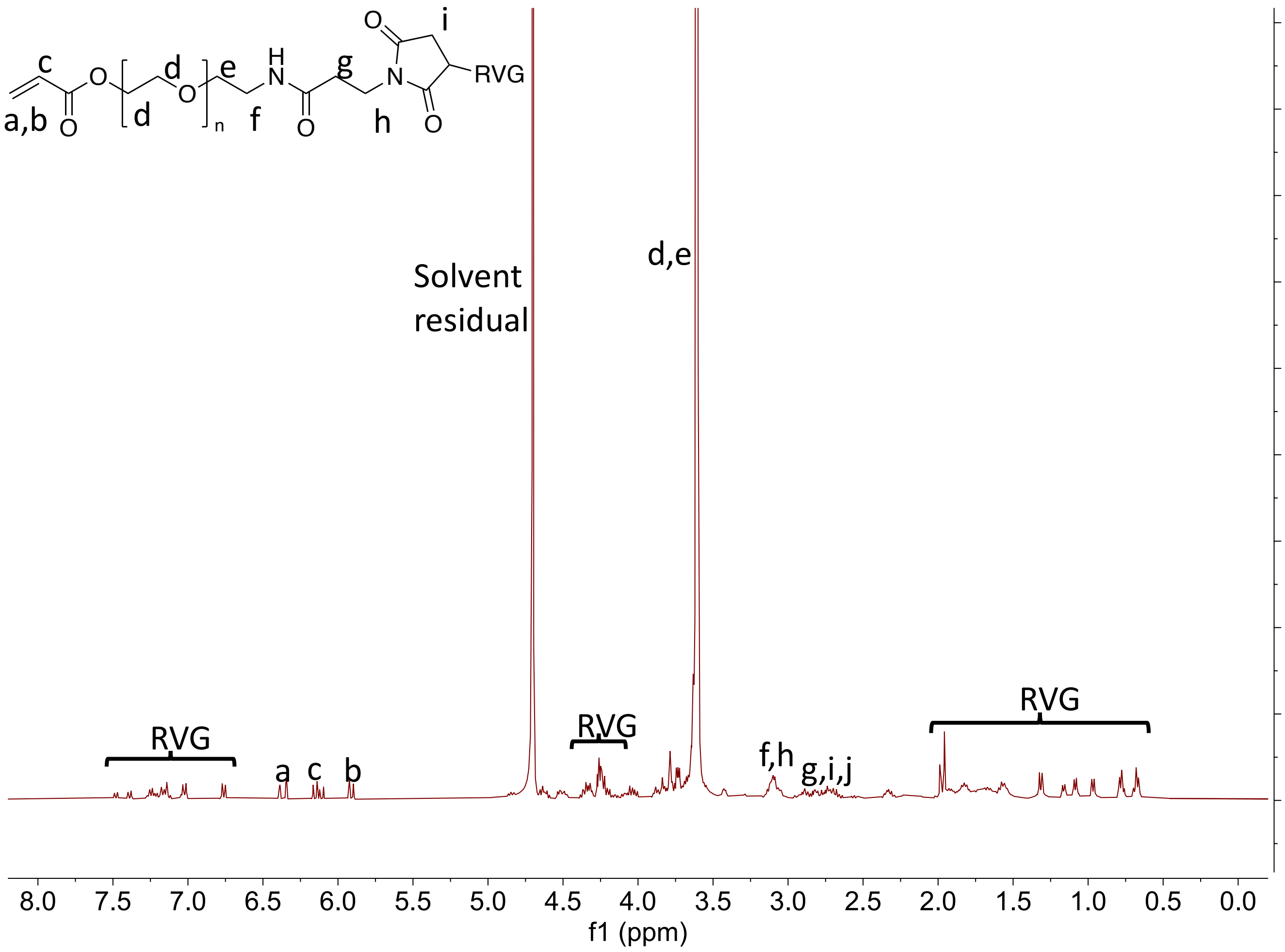


**Supp. Figure 5.** $^{1}$H NMR spectrum of Ac-PEG-RVG

Supplementary Materials

**Supp. Table 1.** Information about the animals evaluated in this project. When possible, graphs in the manuscript use a specific symbol and color to identify which animal corresponds to each data point; the animal corresponding to each symbol and color is specified in this table. In the 'Hemispheres analyzed' column, a "full well of tissue" refers to one sixth serial coronal brain sections spaced 240 µm apart. Storage buffer is defined in the methods section of the manuscript. CPP, cell penetrating peptide; F, female; HE, hematoxylin and eosin; L, left; M, male; NC, nanocapsule; n/a, not applicable; PBS, phosphate buffered saline; R, right; ROI, region of interest; RVG, rabies virus glycoprotein; UW-Madison, University of Wisconsin - Madison.

| Mouse ID | Sex | Age injected (days) | Weight (g) | Facility injected | L hemisphere injection | R hemisphere injection | Volume injected (µl) | Injection rate (µl/min) | Solution NCs suspended in | Hemispheres analyzed | Symbol used in graphs |
|---|---|---|---|---|---|---|---|---|---|---|---|
| J1 | F | 51 | 19.9 | The Jackson Laboratory | NC-RVG, negative control | NC-RVG, Ai14 targeting | 1 | 0.2 | Storage Buffer | R for ROI size and percent editing (full well of tissue); R for types of neurons edited (full well of tissue) | purple circle |
| J2 | F | 58 | 20.6 | The Jackson Laboratory | NC-RVG, negative control | NC-RVG, Ai14 targeting | 1 | 0.2 | Storage Buffer | R for ROI size and percent editing (full well of tissue) | purple square |
| J3 | F | 65 | 17.7 | The Jackson Laboratory | NC-RVG, negative control | NC-RVG, Ai14 targeting | 1 | 0.2 | Storage Buffer | R for ROI size and percent editing (full well of tissue) | purple triangle |
| J4 | M | 58 | 25.2 | The Jackson Laboratory | NC-RVG, negative control | NC-RVG, Ai14 targeting | 1 | 0.2 | Storage Buffer | R for ROI size and percent editing (full well of tissue); R for types of neurons edited (full well of tissue) | purple diamond |
| J5 | M | 58 | 20.1 | The Jackson Laboratory | NC-RVG, negative control | NC-RVG, Ai14 targeting | 1 | 0.2 | Storage Buffer | R for ROI size and percent editing (full well of tissue); R for types of neurons edited (full well of tissue) | purple open circle |
| J6 | M | 58 | 23.9 | The Jackson Laboratory | NC-RVG, negative control | NC-RVG, Ai14 targeting | 1 | 0.2 | Storage Buffer | R for ROI size and percent editing (full well of tissue) | purple open square |
| J7 | F | 51 | 19.3 | The Jackson Laboratory | NC-no ligand, negative control | NC-no ligand, Ai14 targeting | 1 | 0.2 | Storage Buffer | R for ROI size and percent editing (full well of tissue); R & L for HE (full well of tissue); R & L for glial cells (full well of tissue) | red circle |
| J8 | F | 58 | 17.9 | The Jackson Laboratory | NC-no ligand, negative control | NC-no ligand, Ai14 targeting | 1 | 0.2 | Storage Buffer | R for ROI size and percent editing (full well of tissue); R & L for HE (full well of tissue); R & L for glial cells (full well of tissue); R for types of neurons edtied (full well of tissue) | red square |

| | | | | | | | | | | | |
|---|---|---|---|---|---|---|---|---|---|---|---|
| J9 | F | 65 | 19.5 | The Jackson Laboratory | NC-no ligand, negative control | NC-no ligand, Ai14 targeting | 1 | 0.2 | Storage Buffer | R for ROI size and percent editing (full well of tissue); R for types of neurons edited (full well of tissue) | red triangle |
| J10 | M | 59 | 19.8 | The Jackson Laboratory | NC-no ligand, negative control | NC-no ligand, Ai14 targeting | 1 | 0.2 | Storage Buffer | R for ROI size and percent editing (full well of tissue) | red diamond |
| J11 | M | 59 | 19.4 | The Jackson Laboratory | NC-no ligand, negative control | NC-no ligand, Ai14 targeting | 1 | 0.2 | Storage Buffer | R for ROI size and percent editing (full well of tissue) | red open circle |
| J12 | M | 59 | 23.6 | The Jackson Laboratory | NC-no ligand, negative control | NC-no ligand, Ai14 targeting | 1 | 0.2 | Storage Buffer | R for ROI size and percent editing (full well of tissue); R & L for HE (full well of tissue); R & L for glial cells (full well of tissue); R for types of neurons edited (full well of tissue) | red open square |
| J13 | F | 52 | 18.4 | The Jackson Laboratory | NC-CPP, negative control | NC-CPP, Ai14 targeting | 1 | 0.2 | Storage Buffer | R for ROI size and percent editing (full well of tissue); R for types of neurons edited (full well of tissue) | blue circle |
| J14 | F | 59 | 18.8 | The Jackson Laboratory | NC-CPP, negative control | NC-CPP, Ai14 targeting | 1 | 0.2 | Storage Buffer | R for ROI size and percent editing (full well of tissue); R for types of neurons edited (full well of tissue) | blue square |
| J15 | F | 66 | 18.3 | The Jackson Laboratory | NC-CPP, negative control | NC-CPP, Ai14 targeting | 1 | 0.2 | Storage Buffer | R for ROI size and percent editing (full well of tissue) | blue triangle |
| J16 | M | 59 | 23.3 | The Jackson Laboratory | NC-CPP, negative control | NC-CPP, Ai14 targeting | 1 | 0.2 | Storage Buffer | R for ROI size and percent editing (full well of tissue) | blue diamond |
| J17 | M | 59 | 25.5 | The Jackson Laboratory | NC-CPP, negative control | NC-CPP, Ai14 targeting | 1 | 0.2 | Storage Buffer | R for ROI size and percent editing (full well of tissue); R for types of neurons edited (full well of tissue) | blue open circle |
| J18 | M | 59 | 19.8 | The Jackson Laboratory | NC-CPP, negative control | NC-CPP, Ai14 targeting | 1 | 0.2 | Storage Buffer | R for ROI size and percent editing (full well of tissue) | blue open square |
| UW1 | F | 129 | 21 | UW-Madison | NC-CPP, Ai14 targeting | NC-CPP, negative control | 1.5 | 0.2 | PBS | L for ROI size and percent edited (full well of tissue) | blue circle - Supp. Fig 1 |
| UW2 | F | 129 | 23 | UW-Madison | NC-no ligand, Ai14 targeting | NC-no ligand, negative control | 1.5 | 0.2 | PBS | L for ROI size and percent edited (full well of tissue) | red square - Supp. Fig. 1 |

| | | | | | | | | | | | |
|---|---|---|---|---|---|---|---|---|---|---|---|
| UW3 | F | 125 | 21 | UW-Madison | NC-no ligand, Ai14 targeting | NC-no ligand, negative control | 1.5 | 0.2 | PBS | L for ROI size and percent edited (full well of tissue) | red circle - Supp. Fig. 1 |
| UW4 | M | 83 | 32 | UW-Madison | NC-CPP, Ai14 targeting | NC-CPP, Ai14 targeting | 1.5 | 0.2 | PBS | R for ROI size and percent editing (full well of tissue) | blue triangle - Supp. Fig. 1 |
| UW5 | M | 83 | 28 | UW-Madison | NC-CPP, Ai14 targeting | NC-CPP, Ai14 targeting | 1.5 | 0.2 | PBS | R for ROI size and percent editing (full well of tissue) | blue square - Supp. Fig. 1 |
| UW6 | F | 96 | 21 | UW-Madison | NC-no ligand, Ai14 targeting | NC-no ligand, Ai14 targeting | 1.5 | 0.2 | PBS | R and L for ROI size and percent editing (full well of tissue) | red triangle L hemisphere; red diamond R hemisphere - Supp. Fig. 1 |
| UW7 | F | 90 | 23 | UW-Madison | Uninjected | n/a | n/a | n/a | n/a | L side for glial cells (full well of tissue) | black circle |
| UW8 | F | 90 | 21 | UW-Madison | Uninjected | n/a | n/a | n/a | n/a | L side for glial cells (full well of tissue); L side for HE | black square |
| UW9 | F | 88 | 24 | UW-Madison | Uninjected | n/a | n/a | n/a | n/a | L side for glial cells (full well of tissue) | black triangle |

Supplementary Materials

**Supp. Table 2.** Primary antibody information

| Goal of the immunostain | Protein labeled | Company | Species | Catalog # | Lot # | Antibody registry # | Dilution | Blocking agent |
|---|---|---|---|---|---|---|---|---|
| Identify neurons and astrocytes expressing TdTomato | Red fluorescent protein (tdTomato) | Rockland | Rabbit | 600-401-379 | 42872 | *AB_2209751* | 1:5000 | Normal Serum |
| | Glial fibrillary acidic protein (GFAP) | Invitrogen | Rat | 13-0300 | VB298933 | *AB_86543* | 1:500 | Normal Serum |
| | Neuronal nuclear protein (NeuN) | Abcam | Mouse | Ab104224 | GR3298963-3 | *AB_10711040* | 1:1000 | Normal Serum |
| Identify gliosis (astrocytes and microglia) | Red fluorescent protein (tdTomato) | Rockland | Rabbit | 600-401-379 | 42872 | *AB_2209751* | 1:5000 | Normal Serum |
| | Glial fibrillary acidic protein (GFAP) | Invitrogen | Rat | 13-0300 | VB298933 | *AB_86543* | 1:500 | Normal Serum |
| | Ionized calcium binding adaptor molecule 1 (Iba1) | Abcam | Goat | Ab5076 | GR3190885-4 | *AB_91676* | 1:100 | Normal Serum |
| Identify type of neurons genome edited (medium spiny neurons or cholinergic interneurons) | Red fluorescent protein (tdTomato) | Rockland | Rabbit | 600-401-379 | 42872 | *AB_2209751* | 1:5000 | Normal Serum |
| | Dopamine- and cyclic-AMP-regulated phosphoprotein of molecular weight 32 kDa (DARPP32) | Santa Cruz Biotechnology | Mouse | sc-271111 | I1021 | AB_10610055 | 1:200 | Normal Serum |
| | Choline acetyltransferase (ChAT) | Novus Biologicals | Goat | NBP1-30052 | LH321a | AB_1968484 | 1:500 | Normal Serum |
| Identify type of neurons genome edited (parvalbumin or calretinin interneurons) | Red fluorescent protein (tdTomato) | Rockland | Rabbit | 600-401-379 | 42872 | *AB_2209751* | 1:5000 | Normal Serum |
| | Parvalbumin | Synaptic Systems | Guinea Pig | 195004 | 3-38 | AB_2156476 | 1:1000 | Normal Serum |
| | Calretinin | R&D Systems | Goat | AF5065 | CBJR0221031 | AB_2068516 | 1:1000 | Normal Serum |

Supplementary Materials

**Supp. Table 3.** Secondary antibody information

| **Primary this secondary was used with** | **Species** | **Catalog #** | **Lot #** | **Dilution** | **Blocking agent** |
|---|---|---|---|---|---|
| Red flourescent protein (tdTomato) | Donkey Anti-Rabbit | A21207 | 1668652 | 1:1000 | Blocking Solution |
| Glial fibrillary acidic protein (GFAP) | Donkey Anti-Rat | A21208 | 2063330 | 1:1000 | Blocking Solution |
| Neuronal nuclear protein (NeuN) | Donkey Anti-Mouse | A21202 | 2147618 | 1:1000 | Blocking Solution |
| Ionized calcium binding adaptor molecule 1 (Iba1) | Donkey Anti-Goat | Ab150135 | GR3324428-3 | 1:1000 | Blocking Solution |
| Choline acetyltransferase (ChAT) | Donkey Anti-Goat | Ab150129 | GR3375501-2 | 1:1000 | Blocking Solution |
| Dopamine- and cyclic-AMP-regulated phosphoprotein of molecular weight 32 kDa (DARPP32) | Donkey Anti-Mouse | AB15107 | GR3234832-4 | 1:1000 | Blocking Solution |
| Parvalbumin | Donkey Anti-Guinea Pig | 7065454148 | 158591 | 1:1000 | Blocking Solution |
| Calretinin | Donkey Anti-Goat | ab150135 | GR3324428-3 | 1:1000 | Blocking Solution |

**Supp. Table 4**. Information on adjustments to LUTs for images used in manuscript figures. Note that some images were acquired at UW-Madison and some at The Jackson Laboratory, and the full range of the LUT scale is different for these images (The Jackson Laboratory full range 16bit: 0,65535; UW-Madison full range 12bit: 0,4095). More information on the microscopes used at either location can be found in the methods. Images as they are shown in manuscript figures sometimes show certain channels in different colors than typically used for each laser - for example, in the tdTomato/NeuN/GFAP stains, GFAP was imaged with a far-red laser and shown in the figures as white. The original lasers used for each image acquisition are clarified below so that LUTs in the table refer to these original lasers used. LUT, look up table; GFAP, glial fibrillary acidic protein; NeuN, neuronal nuclear protein; Iba, ionized calcium binding adaptor molecule 1; DAPI, 4′,6-diamidino-2-phenylindole; DARPP32, dopamine- and cyclic-AMP-regulated phosphoprotein of molecular weight 32kDA; ChAT, choline acetyltransferase.

| Figure number | Figure panel | Location image acquired | Objective image acquired | Original lasers used | 405 LUTs | 488 LUTs | 561 LUTs | 640 LUTs |
|---|---|---|---|---|---|---|---|---|
| 2 | d | The Jackson Laboratory | 20x | 405 = DAPI; 488 = NeuN; 561 = tdTomato; 640 = GFAP | 0, 45000 | 5000, 65535 | 0, 50000 | 15000, 65535 |
| 3 | a | UW-Madison | 4x (whole coronal section) | 405 = DAPI; 488 = NeuN; 561 = tdTomato; 640 = GFAP | 0, 4095 | 0, 3004 | 56, 2067 | 0, 2938 |
| 3 | a | UW-Madison | 40x (higher mag panels) | 405 = DAPI; 488 = NeuN; 561 = tdTomato; 640 = GFAP | 0, 2963 | 195, 2759 | 0, 2890 | 0, 2906 |
| 3 | b | The Jackson Laboratory | 20x | 405 = DAPI; 488 = NeuN; 561 = tdTomato; 640 = GFAP | 0, 45000 | 5000, 65535 | 0, 50000 | 15000, 65535 |
| 4 | a,b,c | UW-Madison | 20x | 405 = DAPI; 488 = ChAT; 561 = tdTomato; 640 = DARPP32 | channel not shown in fig | 500, 1800 | 0, 2000 | Animal J5: 300, 2000; Animal J9: 200, 2000; Animal J17: 300, 3000 |
| 4 | d,e,f | UW-Madison | 20x | 405 = DAPI; 488 = Parvalbumin; 561 = tdTomato; 640 = Calretinin | channel not shown in fig | Animal J8: 0, 994; Animal J13: 0, 2175; Animal J4: 0, 407 | Animal J8: 0, 490; Animal J13: 0, 4095; Animal J4: 0, 290 | Animal J8: 132, 1726; Animal J13: 794, 4095; Animal J4: 75, 1823 |
| 5 | d-i | UW-Madison | 20x | 405 = DAPI; 488 = GFAP; 561 = tdTomato; 640 = Iba1 | 0, 3500 | 200, 3000 | 0, 3500 | 150, 3000 |

| | | | | | | | | |
|---|---|---|---|---|---|---|---|---|
| Supp Fig 1 | a,b,f,g | UW-Madison | 4x | 405 = DAPI; 488 = NeuN; 561 = tdTomato; 640 = GFAP | Animal UW1: 0, 1359; Animal UW2: 0, 4095 | Animal UW1: 0, 1489; Animal UW2: 0, 3394 | Animal UW1: 0, 1025; Animal UW2: 0, 2222 | Animal UW1: 0, 1302; Animal UW2: 0, 2556 |
| Supp Fig 1 | c,d,e,h,I,j | UW-Madison | 20x | 405 = DAPI; 488 = NeuN; 561 = tdTomato; 640 = GFAP | Animal UW1: 0, 3201; Animal UW2: 0, 4095 | Animal UW1: 0, 1994; Animal UW2: 0, 2002 | Animal UW1: 0, 2515; Animal UW2: 0, 2507 | Animal UW1: 0, 2995; Animal UW2: 0, 2995 |
| Supp Fig 2 | a,b,c | UW-Madison | 20x | 405 = DAPI; 488 = ChAT; 561 = tdTomato; 640 = DARPP32 | 0, 4095 | 500, 1800 | 0, 2000 | Animal J5: 300, 2000; Animal J9: 200, 2000; Animal J17: 300, 3000 |
| Supp Fig 3 | a,b,c | UW-Madison | 20x | 405 = DAPI; 488 = Parvalbumin 561 = tdTomato; 640 = Calretinin | Animal J8: 0, 1104; Animal J13: 0, 2430; Animal J4: 0, 580 | Animal J8: 0, 994; Animal J13: 0, 2175; Animal J4: 0, 407 | Animal J8: 0, 490; Animal J13: 0, 4095; Animal J4: 0, 290 | Animal J8: 132, 1726; Animal J13: 794, 4095; Animal J4: 75, 1823 |

**Supp. Table 5.** Comparison of the number of neurons counted using the automated counts in FIJI vs. manual counts. See methods for more details on the counting methods. Automated NeuN counts using this method significantly correlated with manual counts performed in a subset in images ($\rho$ = 0.897, $p$ <0.0001).

| Mouse ID | ROI ID | # Neurons (NeuN+) in ROI counted via FIJI method with watershed correction | # Neurons (NeuN+) in ROI counted via manual counts |
|---|---|---|---|
| J1 | 2_5 | 429 | 448 |
| J1 | 1_1 | 354 | 392 |
| J1 | 1_5 | 346 | 374 |
| J10 | 1_3 | 679 | 659 |
| J10 | 2_3 | 501 | 446 |
| J10 | 1_4 | 367 | 317 |
| J11 | 2_6 | 855 | 449 |
| J12 | 1_3 | 988 | 978 |
| J13 | 1_4 | 486 | 429 |
| J14 | 1_3 | 551 | 585 |
| J15 | 1_6 | 471 | 500 |
| J17 | 1_1 | 413 | 393 |
| J2 | 1_6 | 651 | 668 |
| J4 | 2_3 | 697 | 726 |
| J6 | 1_3 | 224 | 220 |
| J7 | 1_3 | 442 | 375 |
| J8 | 2_3 | 499 | 543 |
| J9 | 2_4 | 352 | 314 |
